# Glycan Reachability Analysis: A Bottleneck-Aware Framework for Inferring Tissue-Specific Glycan Biosynthetic Potential from Transcriptomics

**DOI:** 10.64898/2026.03.24.714093

**Authors:** Yusuke Matsui

## Abstract

Glycan biosynthesis requires the coordinated expression of glycosyltransferases, modifying enzymes, and nucleotide-sugar synthesis and transport machinery. Existing computational tools predict glycan structures from gene expression using binary thresholds, losing quantitative information about relative biosynthetic capacity across tissues. Here we present glycan biosynthetic reachability analysis, which integrates expression-based Z-scores across curated pathway steps using AND/OR logic and minimum aggregation to produce continuous, tissue-comparable scores and an explicit expression-limiting step. Applied to 17,382 GTEx v8 RNA-seq samples across 54 human tissue types, reachability resolved quantitative differences hidden by presence/absence calls; for example, pancreas was 96% binary-positive for the sLeX pathway but had low median reachability (*Z* = −1.86). Bottleneck stability was evaluated across all 19 multi-step metrics. In independent HEK293 knockout glycomics, the minimum score contained information beyond a within-knockout permutation null but did not outperform naive mean aggregation or binary topology. Nested leave-one-knockout-out selection favored a relaxed low quantile (*q* = 0.2) rather than validating the strict minimum. Within-GTEx associations between reachability and signaling-response transcripts are reported only as transcriptomic coherence because predictors and readouts share the same RNA-seq source. The mouse tissue-glycome comparison remained null. Reachability is therefore a hypothesis-generating rank of transcriptomic potential, not a measure of enzymatic activity or glycan abundance.

**Author Summary:** Sugars attached to the surface of every human cell — collectively called glycans — control many biological processes from immune recognition to cancer signaling. Understanding which glycans each tissue can produce requires knowing which sugar-building enzymes are expressed. Current computational approaches often check whether each enzyme is detectable, ignoring quantitative differences in expression. We developed glycan reachability analysis, a method that treats glycan assembly like a production line whose transcriptomic score is set by the weakest expressed step. Using gene expression data from 54 human tissue types, we show that this bottleneck-aware score reveals differences invisible to binary methods; for example, pancreas has all sialyl Lewis X enzymes detectable but at uniformly low transcript levels. In an independent HEK293 knockout-glycomics benchmark, the minimum score contained non-random information but did not outperform naive mean aggregation or binary topology. Associations between reachability and signaling-response transcripts are interpreted only as same-transcriptome coherence and do not establish glycan-mediated signaling. Bottleneck identities were reproducible for most tissue–metric combinations but unstable in some cases, including bulk-brain GM3. Because the mouse tissue-glycome comparison remained null, reachability is presented as a hypothesis-generating rank of transcriptomic potential rather than a substitute for glycomics.

## 1 Introduction

Glycosylation is among the most prevalent and complex post-translational modifications, with the human glycome estimated to encompass thousands of distinct structures synthesized by approximately 700 glycosylation-related genes [1, 2]. The tissue-specific expression of glycosyltransferases, glycosidases, and nucleotide sugar metabolism genes gives rise to organ-specific glycan repertoires that influence cell signaling, immune recognition, and pathogen interactions [3].

Glycan biosynthesis has been modeled with kinetic/ODE systems, Markov chains, and rule-based reaction networks [6, 12-15]; these mechanistic approaches complement transcriptomics-based methods but generally require parameters or reaction rules that are unavailable across many tissues.

Recent advances in computational glycobiology have produced tools that leverage transcriptomic data to predict glycan structures. GlycoMaple [4, 5] maps RNA-seq expression onto 21 curated glycan metabolic pathways, predicting the presence or absence of glycan structures based on whether pathway enzymes exceed a TPM threshold of 1.0. Glycologue [6] uses rule-based logic to predict products from a set of active enzymes. glycoPATH [7] employs machine learning regression to predict quantitative N-glycan profiles, but requires paired glycomic–transcriptomic training data and is limited to N-glycans. Glycopacity [8] and glycoCARTA [9] analyze glycogene expression patterns from single-cell transcriptomics but do not integrate pathway logic to assess biosynthetic potential.

A common limitation of these approaches is that they either (i) reduce expression to binary presence/absence, discarding quantitative information about relative biosynthetic potential, or (ii) require experimental glycomic data for training. None provides a framework for statistically comparing biosynthetic potential across tissues or conditions using expression data alone. Here we address this gap with glycan reachability analysis. We define the reachability of a glycan structure as the minimum normalized expression across all required biosynthetic steps — including both enzymatic reactions and nucleotide sugar donor pathways. This continuous score, termed *biosynthetic reachability* (or simply reachability hereafter), operationalizes a simple heuristic: the least-expressed pathway component constrains the overall transcriptomic potential of the pathway. The underlying assumption is that, when enzymes operate in the linear (unsaturated) regime where reaction rate scales approximately with enzyme concentration, the weakest link in the chain sets an upper bound on pathway output. This assumption is strongest when substrate concentrations are below *K*_*m*_ for most enzymes, a condition that may not hold universally but provides a useful first-order approximation. We emphasize that reachability is a relative rank of transcriptomic biosynthetic potential, not a measure of enzymatic activity or glycan abundance. Its intended use is to prioritize tissue–pathway pairs and candidate limiting steps for direct glycomic measurement; whether the ranking reflects realized glycan abundance must be tested rather than assumed. Unlike metabolic control analysis (MCA), which demonstrates that flux control is distributed among multiple enzymes with flux control coefficients summing to unity [35], our approach does not claim to predict metabolic flux or identify true rate-limiting steps. Rather, it provides a rank-ordering of tissues by their transcriptomic capacity for each biosynthetic pathway.

## 2 Methods

### 2.1 Reachability score definition

For a glycan biosynthetic pathway *P* consisting of enzymatic steps *E*_1_, *E*_2_, … , *E*_*k*_ and required donor substrates *D*_1_, *D*_2_, … , *D*_*m*_, the reachability score for sample *s* is defined as:

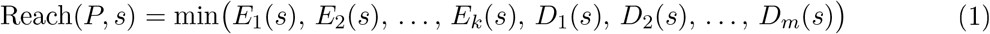

Each enzymatic step *E*_*i*_ may be catalyzed by one or more isozymes. In the primary score, we aggregate isozyme contributions using the arithmetic mean:

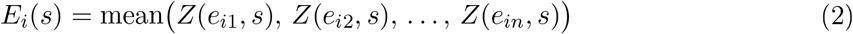

Sensitivity to OR=max and to using ST6GAL1 alone is detailed in Supplementary Methods and Supplementary File S2. Multi-subunit complexes remain AND nodes.

where *Z*(*g, s*) is the per-gene Z-score computed as:

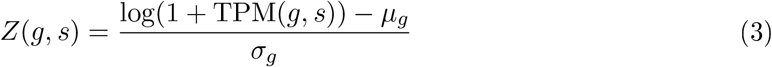

with *µ*_*g*_ and *σ*_*g*_ being the mean and standard deviation of log(1 + TPM) across all samples.

Each donor substrate *D*_*j*_ requires the coordinated expression of synthesis enzymes, activating enzymes, and Golgi transporters with AND logic:

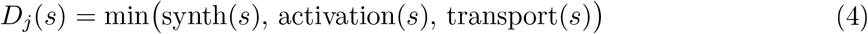

For donor substrates synthesized by alternative pathways (e.g., GDP-Fuc via de novo or salvage), OR logic is applied between pathways:

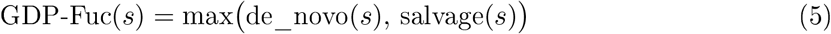

### 2.2 Glycan pathways modeled

Five glycan families were modeled, yielding 23 reachability metrics (Tables 1–2; pathway schematics in Fig 1A–E; metric definitions in Fig 1F; machine-readable definitions in Supplementary File S1). Several biological simplifications are noted: MGAT4A/B (tri-antennary branching) and MGAT5 (tetra-antennary branching) are grouped as a single OR step despite catalyzing distinct reactions; the HS3ST family (HS3ST1–6) is modeled as one OR group despite distinct substrate specificities; and B4GALT1–6 are grouped for LacNAc synthesis despite differential glycoprotein/glycolipid preferences (see Supplementary Methods for sensitivity analyses). Within each pathway, enzymatic steps are connected by AND logic (min-aggregation; all components required). Where multiple isozymes catalyze the same reaction, OR=mean is applied. Enzyme complexes are modeled with appropriate subunit logic: EXT1/EXT2 (required heteromeric complex, AND), ALG13/ALG14 (required heterocomplex, AND), and the OST complex (catalytic STT3A/B as OR; structural subunits RPN1, RPN2, DAD1, DDOST, OSTC as AND; oxidoreductase TUSC3/MAGT1 as OR). All LLO assembly genes (DPAGT1 through ALG10) are modeled as sequential AND steps. Five nucleotide sugar donor supply chains, shared across multiple pathways, are summarized in Table 1. GALE appears in both the UDP-Gal and UDP-GalNAc donor pathways, as it catalyzes epimerization of both UDP-Glc/Gal and UDP-GlcNAc/GalNAc [32].

**Table 1:** Nucleotide sugar donor substrate supply chains. Each uses AND (min) logic across sequential biosynthetic, activation, and Golgi transport steps. GDP-Fuc uniquely uses OR_alt_ (max) between de novo and salvage branches. GALE appears in both UDP-Gal and UDP-GalNAc pathways as it catalyzes both UDP-Glc/Gal and UDP-GlcNAc/GalNAc epimerization.

| Donor | Sequential steps (AND) | Used by |
| --- | --- | --- |
| CMP-Sia | GNE → NANS → NANP → CMAS → SLC35A1 | sLeX, Ganglio., N-gly., O-GalNAc |
| GDP-Fuc | max(GMDS → TSTA3, FUK → FPGT) → SLC35C1 | sLeX, N-glycan |
| UDP-Gal | UGP2 → GALE → SLC35A2 | sLeX, Ganglio., O-GalNAc |
| PAPS | PAPSS1/2 (OR) → SLC35B2/B3 (OR) | HS |
| UDP-GalNAc | UAP1 → GALE → SLC35D1/A2 (OR) | Ganglio., O-GalNAc |

**Table 2:** Glycan biosynthetic pathway steps and reachability metrics. Steps within each pathway are connected by AND (min) logic; isozyme groups use OR=mean. Required donors refer to Table 1. 22 glycan pathway metrics plus PAPS donor supply as a standalone metric (23 total; see S4 Table for a complete listing with pathway family, step count, gene count, and primary/composite classification). SHH-like differs from WNT-like by requiring 3-O sulfation. Display names (Metric column) are used throughout the main text; corresponding code variable names are listed in S4 Table.

| Step | Enzymes (logic) | Metric |
| --- | --- | --- |
| **Sialyl Lewis X (sLeX)** |  | donors: CMP-Sia, GDP-Fuc, UDP-Gal |
| LacNAc formation | B4GALT1–6 (OR) |  |
| α2,3-sialylation | ST3GAL3/4/6 (OR) |  |
| α1,3/4-fucosylation | FUT3/5/6 (OR) | sLeX reachability |
| **Gangliosides** |  | donors: CMP-Sia, UDP-Gal, UDP-GalNAc |
| LacCer synthesis | UGCG + B4GALT5/6 (OR) |  |
| GM3 | + ST3GAL5 | GM3 reach |
| GM2 | + B4GALNT1 | GM2 reach |
| GM1 | + B3GALT4 | GM1 reach |
| GD3 | GM3 + ST8SIA1 | GD3 reach |
| **Heparan sulfate (HS)** |  | donor: PAPS |
| Chain polymerization | EXT1 + EXT2 (AND)* | HS polymerization |
| N-sulfation | NDST1–4 (OR) | HS N-sulfation |
| 2-O-sulfation | HS2ST1 | HS 2-O sulfation |
| 6-O-sulfation | HS6ST1–3 (OR) | HS 6-O sulfation |
| 3-O-sulfation | HS3ST1–6 (OR) | HS 3-O sulfation |
| FGF-like profile | Core + N + 2-O (AND) | HS FGF-like |
| WNT-like profile | Core + N + 6-O (AND) | HS WNT-like |
| SHH-like profile | Core + N + 6-O + 3-O (AND) | HS SHH-like |
| **N-glycan processing** |  | donors: GDP-Fuc (core fuc.), CMP-Sia (sia.) |

Table 2 continued
| Step | Enzymes (logic) | Metric |
| --- | --- | --- |
| LLO initiation | DPAGT1 |  |
| LLO assembly | ALG13/14 (AND)<sup>‡</sup>, ALG1, 2, 11, 3, 9, 12, 6, 8, 10, 5 (AND) |  |
| OST transfer | STT3A/B (OR) + structural** (AND) + TUSC3/MAGT1 (OR) |  |
| Mannosidase I | MAN1A1/A2/B1/C1 (OR) |  |
| GnT-I | MGAT1 |  |
| Mannosidase II | MAN2A1/A2 (OR) |  |
| GnT-II | MGAT2 | Ng complex |
| Branching | MGAT4A/B, MGAT5 (OR) | Ng branch |
| Core fucosylation | FUT8 | Ng core Fuc |
| Bisecting GlcNAc | MGAT3 | Ng bisect |
| Sialylation | ST6GAL1/2 (OR) | Ng sialylation |
| **O-GalNAc (mucin-type)** |  | donors: UDP-GalNAc, UDP-Gal, CMP-Sia |
| Tn initiation | GALNT1–20 (OR) | OGN Tn |
| Core 1 | C1GALT1 + COSMC (AND) | OGN Core 1 |
| Core 2 | GCNT1 | OGN Core 2 |
| Sialylation | ST3GAL1/2 (OR) | OGN sialylation |

\*EXT1/EXT2 form a required heteromeric complex [31]; AND (min) logic. <sup>‡</sup>ALG13/14 form a heterocomplex; all other ALG genes act sequentially (AND); each is essential (CDG). \*\*OST structural subunits: RPN1, RPN2, DAD1, DDOST, OSTC (all required).

**Fig. 1:**
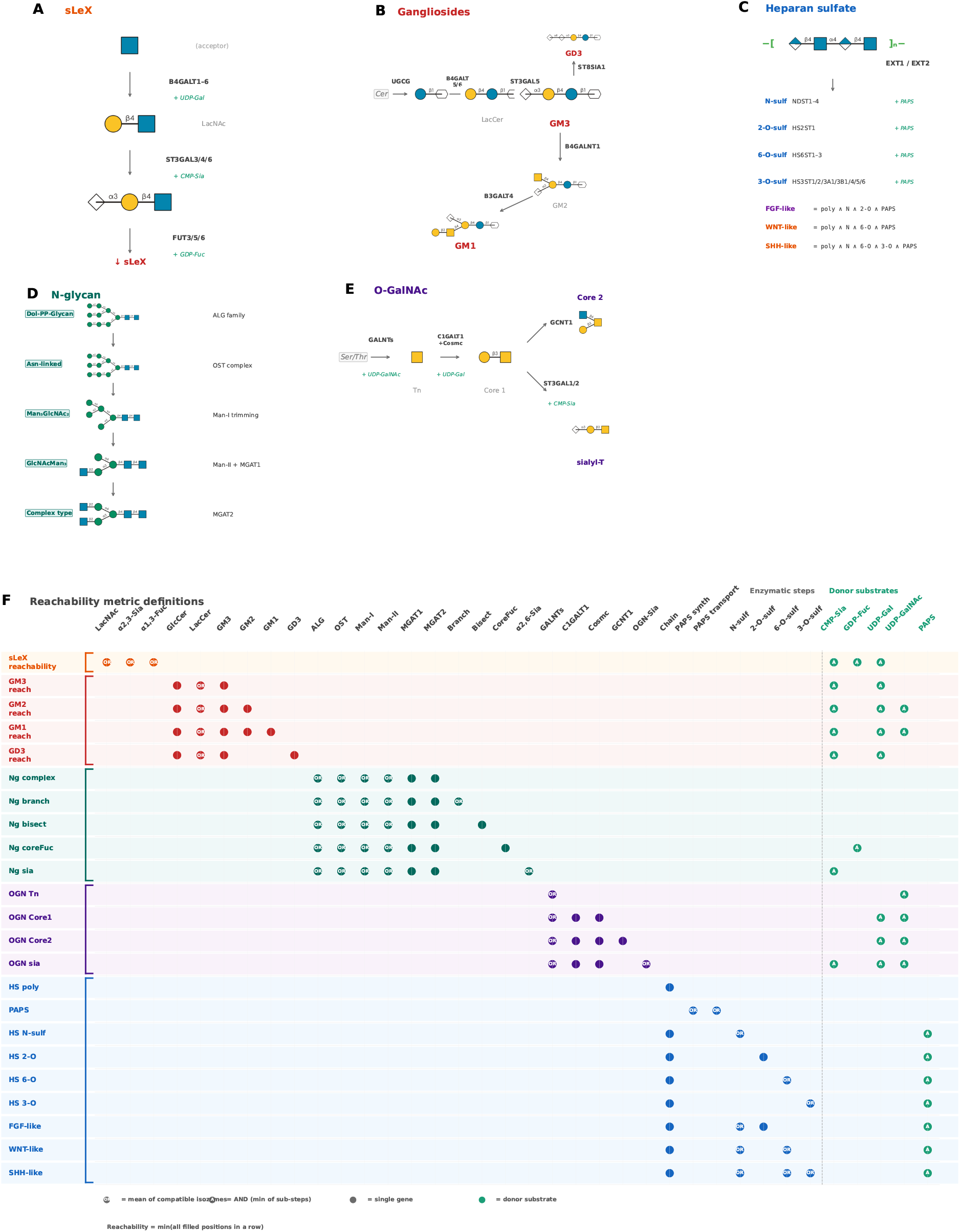
Glycan biosynthesis pathway architecture and reachability metric definitions. **(A-E)** Schematic diagrams of five glycosylation pathway families modeled in this study: (A) sLeX, (B) ganglioside, (C) heparan sulfate, (D) N-glycan, and (E) O-GalNAc. Each diagram shows the sequential enzymatic steps (black) and required nucleotide sugar donors (green). **(F)** Reachability metric definition matrix for all 23 metrics. OR nodes are aggregated by their mean and required sequential steps by their minimum. The score is the minimum across required positions; the argmin is an expression-limiting position, not a proven biochemical rate-limiting step.

### 2.3 Data and preprocessing

GTEx v8 [10]: Gene-level TPM values for 17,382 RNA-seq samples across 54 tissue types from 948 post-mortem donors were obtained from the GTEx Portal (open access; sample sizes per tissue in S3 Table). Gene symbols from the GCT Description field were used; duplicate symbols (multiple Ensembl IDs mapping to the same symbol) were averaged.

### 2.4 Statistical analysis

Kruskal-Wallis tests assessed global tissue-level variation per metric. Pairwise Wilcoxon rank-sum tests with Benjamini–Hochberg (BH) FDR correction were performed for all tissue pairs (1,225 pairs per metric). Clifffs delta was computed as a non-parametric effect size. Effect size guidelines follow Romano et al. (2006): small ≥ 0.147, medium ≥ 0.33, large ≥ 0.474 [34]. Tissues with fewer than 20 samples were excluded from pairwise comparisons (retaining 50 tissues).

The *n* ≥ 20 rule excluded Cervix Ectocervix, Cervix Endocervix, Fallopian Tube, and Kidney Medulla; Bladder (*n* = 21) was retained, yielding 50 tissues. Effect sizes are reported alongside *p*-values.

All *p*-values were corrected using the Benjamini–Hochberg procedure; a detailed multiple testing correction strategy is provided in Supplementary Methods.

### 2.5 Comparison with GlycoMaple-like binary approach

To compare against the threshold-based approach used by GlycoMaple [4], we reimplemented binary pathway assessment using the same TPM ≥ 1 threshold and AND logic described in Huang et al. (2021). For each pathway step, a step was considered “active” if at least one enzyme in the OR group exceeded the threshold. A sample was classified as “reachable” (binary) if all steps were active. Tissue-level binary reachability was computed as the percentage of samples classified as reachable. Spearman correlation between binary percentage and median reachability Z-score was computed across tissues.

### 2.6 Aging analysis

Age-related changes in reachability were assessed using Spearman correlation between donor age (bracket midpoint) and per-sample reachability scores within each tissue. Tissues with fewer than 3 age brackets containing at least 10 samples each were excluded. Correlations were classified as significant decline or increase at FDR *<* 0.05 (BH correction within each tissue across all 23 metrics). Tissue-cascade coherence between age-reachability and age-downstream target correlations was assessed as described in Section 2.7, Panel C.

### 2.7 Transcriptomic coherence between reachability and pathway-response genes

To assess whether reachability is transcriptionally coherent with the response programs of pathways in which these glycans act, we examined three independent glycan-signaling cascades where the glycan product has a well-characterized effect on a receptor signaling pathway. We stress at the outset that this is a coherence analysis *within a single RNA-seq dataset*: both the reachability scores and the pathway-response readouts are derived from the same GTEx transcriptomes, so it cannot, and does not, establish that the glycan products mediate these associations — only that reachability co-varies across tissues with the relevant transcriptional programs. The external test of whether reachability tracks measured glycans is provided separately (Sections 3.9-3.10).

These three cascades were selected *a priori* based on two criteria: (i) the glycan product must have a well-established, experimentally validated effect on a specific receptor signaling pathway with published mechanistic evidence, and (ii) the downstream signaling targets must be canonical transcriptional readouts that are widely used as pathway activity indicators in the literature. Other glycan-signaling relationships exist (e.g., Notch/O-fucosylation, TGF-*β*/HS) but were excluded because their downstream transcriptional signatures are less specific or require cell-type-specific interpretation that bulk tissue RNA-seq cannot resolve. The selected cascades were as follows:

#### (i) WNT cascade

HS N- and 6-O-sulfation (HS WNT-like reachability) promotes WNT ligand binding and co-receptor engagement. Downstream targets: AXIN2, LEF1, TCF7L2, MYC, CCND1, canonical WNT/*β*-catenin transcriptional readouts [16, 17].

#### (ii) EGFR cascade

GM3 ganglioside (GM3 reachability) inhibits EGFR dimerization and activation [18, 19]. Downstream targets: AREG, EREG, EGR1 (direct EGFR-induced transcription), DUSP6, SPRY2 (EGFR/MAPK negative feedback genes widely used as pathway activity indicators [20]). Because GM3 is proposed to inhibit EGFR signaling, the EGFR response score was multiplied by −1 in all coherence analyses. Under this direction-oriented convention, a positive *ρ* would match the proposed inhibition, whereas a negative *ρ* means that the raw GM3-reachability/EGFR-target association is positive and therefore opposite the prespecified direction.

#### (iii) Selectin cascade

sLeX (sLeX reachability) is the minimal glycan determinant for selectin-mediated leukocyte rolling. Downstream targets: ICAM1, VCAM1, ITGB2 (endothelial adhesion molecules engaged after selectin-mediated rolling), SELPLG/PSGL-1 (the primary P-selectin glycoprotein ligand that presents sLeX on its surface; note that SELPLG is a downstream target presenting the glycan, not a biosynthetic enzyme), CCL2, CXCL8 (chemokines induced by selectin-mediated NF-*κ*B activation [21]).

For each cascade, reachability was compared with (i) the mean Z-score of all pathway genes and (ii) the mean Z-score of core catalytic genes. Panel A describes marginal correlations; Panel B describes the conditional association after controlling for the core mean. Neither is interpreted as predictive validation. Core sets were WNT: EXT1/2, NDST1–4, HS6ST1–3; EGFR: UGCG, B4GALT5/6, ST3GAL5; and Selectin: B4GALT1-6, ST3GAL3/4/6, FUT3/5/6.

Analyses were performed at the tissue level using median values across samples within each tissue (*n* = 50 tissues with at least 20 samples).

#### Panel A (tissue-level correlation)

Spearman correlations between each predictor (reachability or naive mean expression) and response score were computed independently for each cascade. The EGFR response was sign-oriented as described above.

#### Panel B (partial correlation)

Spearman partial correlation (`ppcor` [22]) assessed the conditional rank association of reachability and the response score after controlling for core mean expression. A nested linear-model F-test reported incremental *R*^2^ as a complementary descriptive analysis.

#### Panel C (aging cascade coherence)

For each tissue-cascade pair, Spearman correlations of age (bracket midpoint) with reachability and with downstream score were computed at the individual sample level. Tissue-cascade pairs with fewer than 3 age brackets containing at least 10 samples were excluded, yielding 144 evaluable pairs. Concordance between age-reachability and age-downstream correlations was assessed across tissue-cascade pairs.

#### Multiple testing correction

All *p*-values within each panel were adjusted using the Benjamini–Hochberg procedure. Panel A: 6 tests (3 cascades *×* 2 predictors). Panel B: 3 tests per test type (Spearman partial correlation and F-test corrected separately). Panel C: 144 tests per correlation type.

Additional coherence sensitivities and the independent KO-glycomics benchmark are described in Supplementary Methods.

## 3 Results

### 3.1 Tissue-specific glycogene expression recapitulates established programs and motivates a bottleneck-aware integration

To establish the transcriptomic landscape underlying tissue-specific glycan biosynthesis, we surveyed the expression of 97 glycosylation-related genes — spanning 28 functional categories including donor substrate synthesis/transport, glycosyltransferases, glycosidases, and sulfotransferases — across all 54 GTEx tissue types (Fig 1; S1 Fig; S2 Fig). That glycosyltransferase and glycogene expression varies markedly across tissues is itself well established: large-scale transcriptomic surveys have documented these organ-specific programs and their comparatively uniform donor-substrate machinery [8, 9, 11]. We therefore present this landscape not as a novel finding but as the premise for our method — confirming that GTEx recapitulates the known organization before asking whether quantitative pathway integration adds resolution beyond a gene-by-gene view. The contribution of this study is not the expression landscape but the bottleneck-aware, continuous reachability integration that follows.

Hierarchical clustering of median Z-scores at the pathway-category level revealed pronounced tissue-specific expression programs (S1 Fig A). Brain regions formed a tight cluster characterized by elevated ganglioside synthase expression but reduced donor substrate and HS-related gene expression. Connective tissue-rich organs (nerve, fibroblasts, ovary) showed the highest HS-related expression. Gastrointestinal tissues clustered together with elevated sLeX pathway components, consistent with mucosal glycan biology. Whole blood, skeletal muscle, and heart consistently ranked lowest across most categories, reflecting the reduced glycosylation machinery in terminally differentiated cells.

This landscape provides the biological premise for our central question: can quantitative integration of these expression patterns infer tissue-specific glycan biosynthetic potential more accurately than binary threshold approaches?

### 3.2 Reachability analysis: A bottleneck-aware quantitative framework

We developed glycan reachability analysis by integrating per-gene Z-scores with OR=mean for isozyme groups and AND=min across required enzymatic and donor-supply steps (Fig 1F). The minimum is an interpretable expression heuristic; it is not a biochemical flux model, and its comparative performance is evaluated rather than assumed below.

We computed 23 reachability metrics spanning five glycan families for all 17,382 GTEx samples. All metrics showed highly significant tissue-level variation (Kruskal-Wallis *p <* 10^−300^ for all; Table 3). For the 14 primary metrics (sLeX, gangliosides including GM2, and HS), chi-squared statistics ranged from 9,458 (GD3) to 14,649 (HS polymerization). We note that reachability scores are not directly comparable across pathways of different lengths, because pathways with more steps produce systematically lower minimum Z-scores (expected minimum 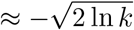 for *k* steps); all cross-tissue comparisons in this study are therefore performed within each metric.

Hierarchical clustering of tissue median reachability scores (S10 Fig) revealed biologically coherent groupings that differed from and complemented the raw expression landscape in S1 Fig A. While the expression landscape shows which genes are transcribed, the reachability heatmap reveals which biosynthetic pathways have expression-complete profiles. For example, brain regions express ganglioside synthases at high levels (S1 Fig A) but show only moderate ganglioside reachability (S10 Fig), because the bottleneck principle identifies donor substrate availability as the potential rate-limiting factor (see Section 3.4).

### 3.3 Reachability provides quantitative resolution complementary to binary approaches

Binary threshold methods such as GlycoMaple [4] address a fundamentally different question — “can this tissue produce glycan X?” — whereas reachability analysis asks “how does this tissuefs transcriptomic potential compare to other tissues?” These approaches are complementary, not competitive, and we compare them here to illustrate the additional information provided by quantitative scoring.

We compared reachability scores against a GlycoMaple-like binary threshold approach (TPM ≥ 1 per enzyme) across 50 tissues with sufficient sample size (*n* ≥ 20).

For sLeX, binary positive rates ranged from *<* 1% (whole blood, skeletal muscle) to 100% (minor salivary gland, esophagus mucosa), while reachability Z-scores ranged from −2.56 to −0.11. The two measures showed a weak, nominal positive correlation (Spearman *ρ* = 0.283, *p* = 0.046; Fig 2A), indicating that binary detection and the quantitative score are related but not interchangeable.

**Fig. 2:**
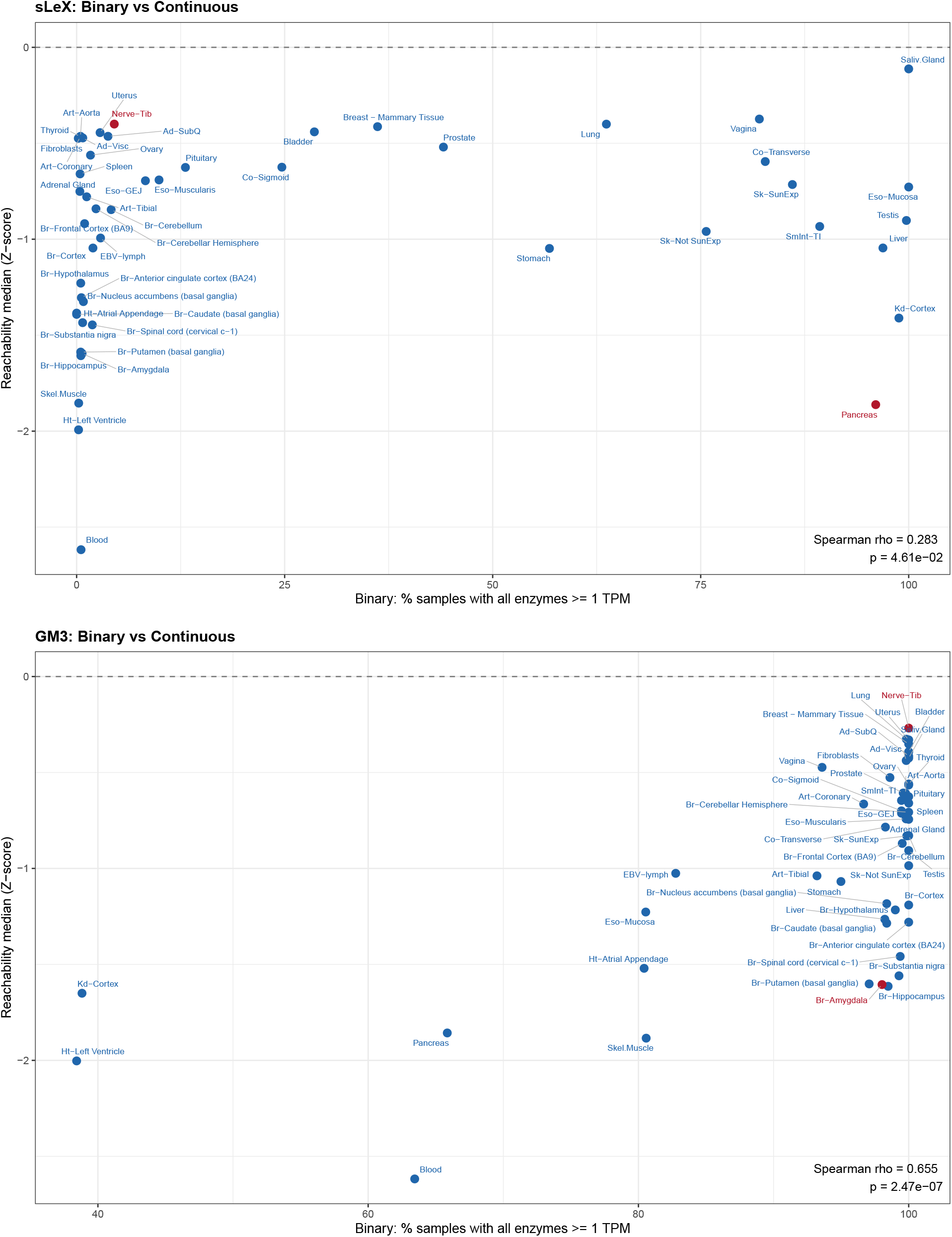
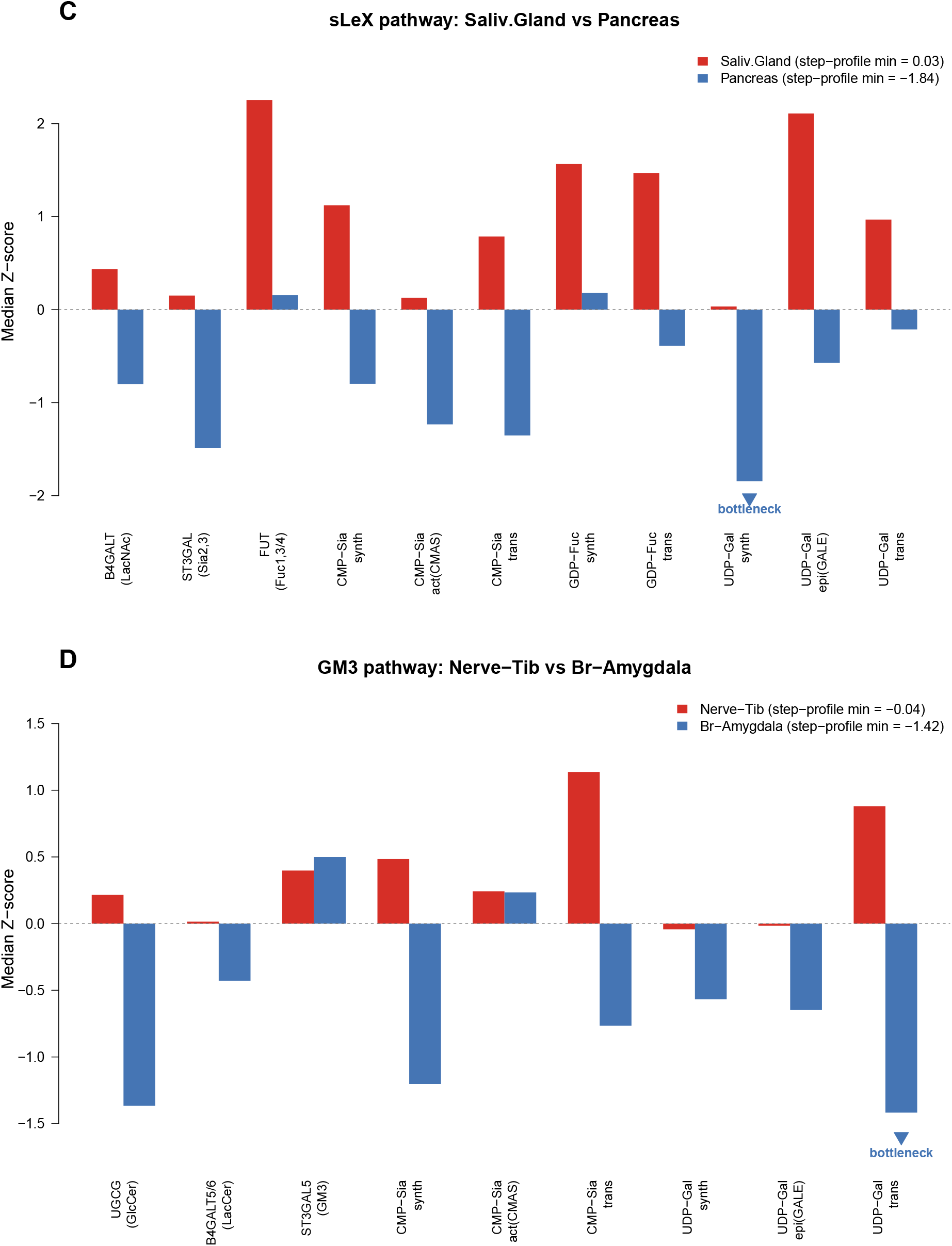
Binary threshold approaches lose quantitative resolution in biosynthetic potential assessment. **(A)** GlycoMaple-like binary-positive rate versus median sLeX reachability across 50 tissues (*ρ* = 0.283, *p* = 0.046). **(B)** Same for GM3 (*ρ* = 0.655, *p* = 2.47 *×* 10^−7^). Discordance describes different score behavior and does not determine which better reflects glycan abundance.**(C)** Per-step median Z-scores for the sLeX pathway comparing minor salivary gland (red) and pancreas (blue). Arrowhead marks the bottleneck in pancreas. **(D)** Per-step median Z-scores for the GM3 pathway comparing nerve-tibial (red) and brain-amygdala (blue). alues labeled “step-profile min” are minima of the displayed tissue-median step profiles; Results Section 3.3 reports medians of per-sample reachability scores. Stability statistics for these argmin calls are provided in Supplementary File S2.

The most striking discordance was observed in pancreas: 96% of samples had all sLeX pathway enzymes above the 1 TPM threshold (binary: “capable”), yet the median reachability Z-score was −1.86 (among the lowest of all tissues). This occurs because each enzyme is individually detectable but expressed at uniformly low levels; the min-aggregation across all steps amplifies these individually modest deficits into a substantially reduced biosynthetic potential score.

We present this discordance as an illustration of the resolution reachability adds over a binary call, not as evidence that reachability is correct here: from transcriptomics alone we cannot determine which representation better reflects the produced sLeX glycome, and matched normal-pancreas glycomics would be required to adjudicate. (The clinical sLeX-related biomarker CA19-9/sialyl Lewis A is a marker of *malignant* pancreatic tissue [38] and speaks to tumour biology, not to the transcriptomic bottleneck of normal pancreas, so we do not invoke it as support.) This case also illustrates a key limitation of bulk tissue analysis: pancreatic ductal cells, although a minority population (*<* 5% of pancreatic mass), are known to express sLeX at higher levels than acinar cells, and their signal is likely masked in bulk RNA-seq data dominated by acinar transcripts. The low reachability score thus reflects the dominant cell typefs program rather than the tissuefs full biosynthetic repertoire.

Conversely, nerve tibial tissue showed only ∼5% binary positive rate for sLeX (due to low FUT3/5/6 expression below threshold), yet its reachability score (−0.40) ranked among the highest, driven by strong expression of upstream pathway components and donor substrate machinery.

For GM3, the binary-continuous correlation was stronger (*ρ* = 0.655, *p* = 2.47 *×* 10^−7^; Fig 2B). Six brain regions nevertheless showed *>* 97% binary positivity and bottom-quartile reachability. This discordance identifies the expression positions driving the score but does not establish that brain ganglioside abundance is low.

### 3.4 Identification of pathway bottlenecks

A key advantage of reachability analysis is the ability to identify which pathway step limits overall biosynthetic potential — information that is entirely absent from binary classifications.

#### sLeX bottleneck analysis (Fig 2C)

We compared the per-step Z-scores between minor salivary gland (highest sLeX reachability, *Z* = −0.11) and pancreas (near-lowest, *Z* = −1.86). Pancreas showed low values across the 11 positions, with the deepest tissue-median deficit in UDP-Gal synthesis (*Z* = −1.84). At the sample level, UDP-Gal was the modal limiting step in 81.7% of pancreas samples. It was recovered in all 1,000 donor bootstrap resamples and both donor halves agreed in all 1,000 random splits (Supplementary File S2). Thus this expression-level argmin is stable, although its relation to produced sLeX still requires glycomics.

#### GM3 bottleneck analysis (Fig 2D)

We compared brain amygdala (98% binary-positive; reachability *Z* = −1.61) with tibial nerve (100%; *Z* = −0.27). ST3GAL5 was similar (brain *Z* = 0.50 vs. nerve *Z* = 0.40), whereas several upstream positions were lower in amygdala, including UGCG (−1.37 vs. 0.22) and UDP-Gal transport (−1.42 vs. 0.88). The tissue-median minimum was UDP-Gal transport, and the sample-level modal group was UDP-Gal; this avoids attributing a heterogeneous tissue to one gene.

#### Failure mode — cell-type heterogeneity

The brain is the most ganglioside-rich organ [37], so the low bulk-brain score is counter-evidence to interpreting reachability as realized abundance. The amygdala modal step occurred in only 38.8% of samples, with donor-half agreement 0.573 and bootstrap recovery 0.825; the bottleneck identity is therefore uncertain. Across all 19 multi-step metrics and 50 tissues, the median modal frequency was 0.608, mean donor-half agreement 0.917, and mean bootstrap modal recovery 0.958. We report these quantities as reliability flags and down-weight diffuse, high-entropy calls. Single-cell or spatial data would be required to separate neuronal and glial programs.

### 3.5 Statistical framework: Quantifying tissue differences

To provide a rigorous statistical foundation for tissue-level comparisons, we performed pairwise Wilcoxon rank-sum tests across all 1,225 tissue pairs for each reachability metric.

Signed effect size heatmaps (Fig 3A,B) display − log_10_(FDR) values signed by the direction of difference for sLeX reachability and HS FGF-like reachability, respectively. These reveal the global pattern of tissue differentiation: for sLeX (Fig 3A), gastrointestinal and salivary tissues form a high-reachability block sharply separated from brain, muscle, and blood. For HS FGF-like (Fig 3B), the pattern inverts, with connective tissue-rich organs showing the highest potential and blood/muscle remaining low.

**Fig. 3:**
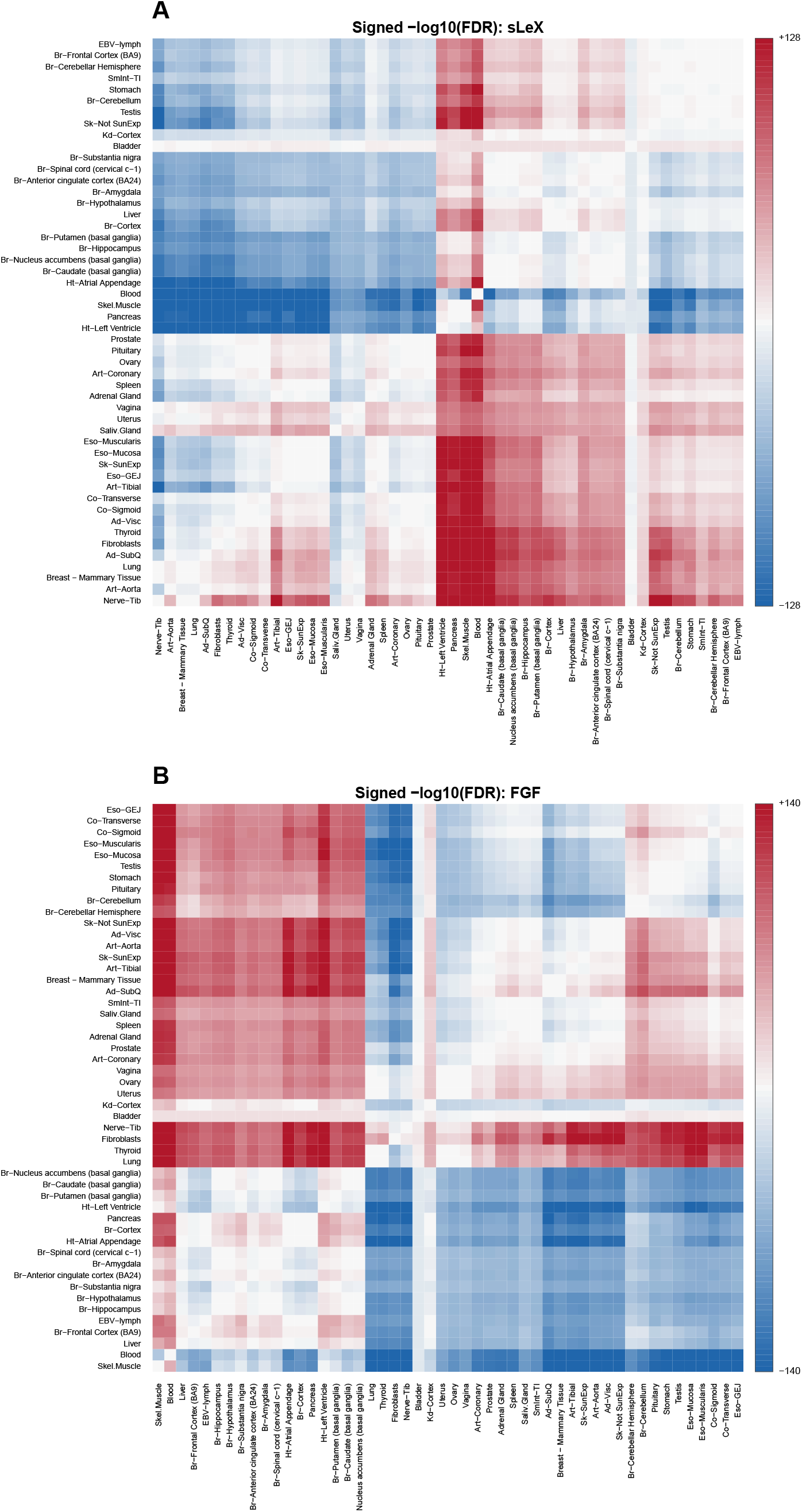
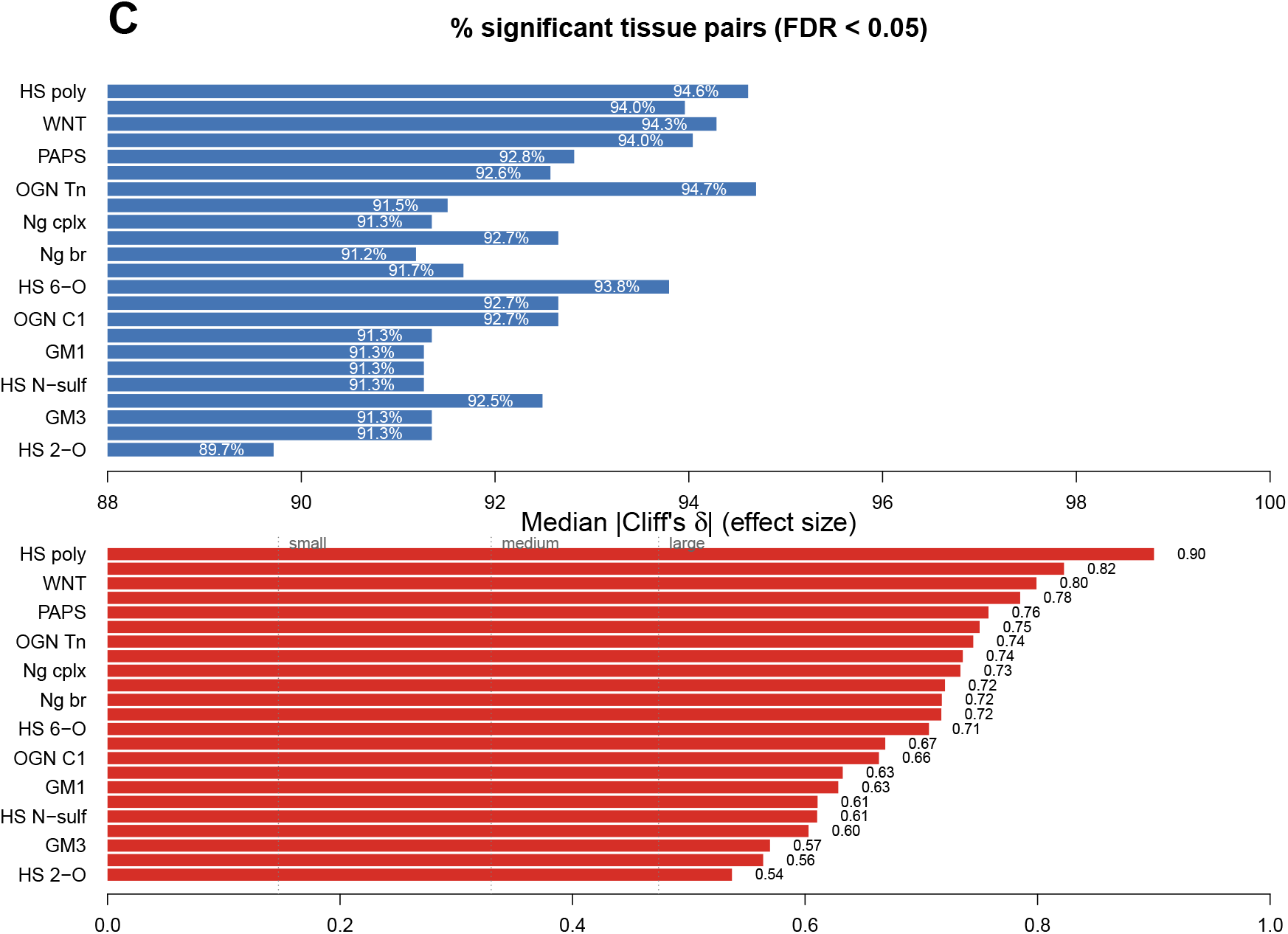
Statistical framework for tissue-level comparison. **(A)** Signed − log_10_(FDR) heatmap for pairwise Wilcoxon tests of sLeX reachability across 50 tissues. **(B)** Same as (A) for HS FGF-like reachability, showing inverted tissue hierarchy compared to sLeX.**(C)** Within-metric FDR-significant proportions and median absolute Clifffs delta for all 23 metrics. In total, 26,025/28,175 comparisons were significant; per-metric proportions were 89.7-94.6%.

Across all 23 metrics, 26,025 of 28,175 tissue-pair tests (92.4%) were significant at within-metric FDR *<* 0.05. Per-metric proportions ranged from 89.7% (HS 2-O sulfation) to 94.6% (HS polymerization), and median absolute Clifffs delta ranged from 0.537 to 0.900 (Fig 3C). These are descriptive transcriptomic differences; their biological consequence is not established by the tests.

### 3.6 Age-related changes in glycan biosynthetic potential

To assess whether glycan biosynthetic potential changes with aging, we computed Spearman correlations between donor age and per-sample reachability scores within each tissue across all 23 metrics (Fig 4).

**Fig. 4:**
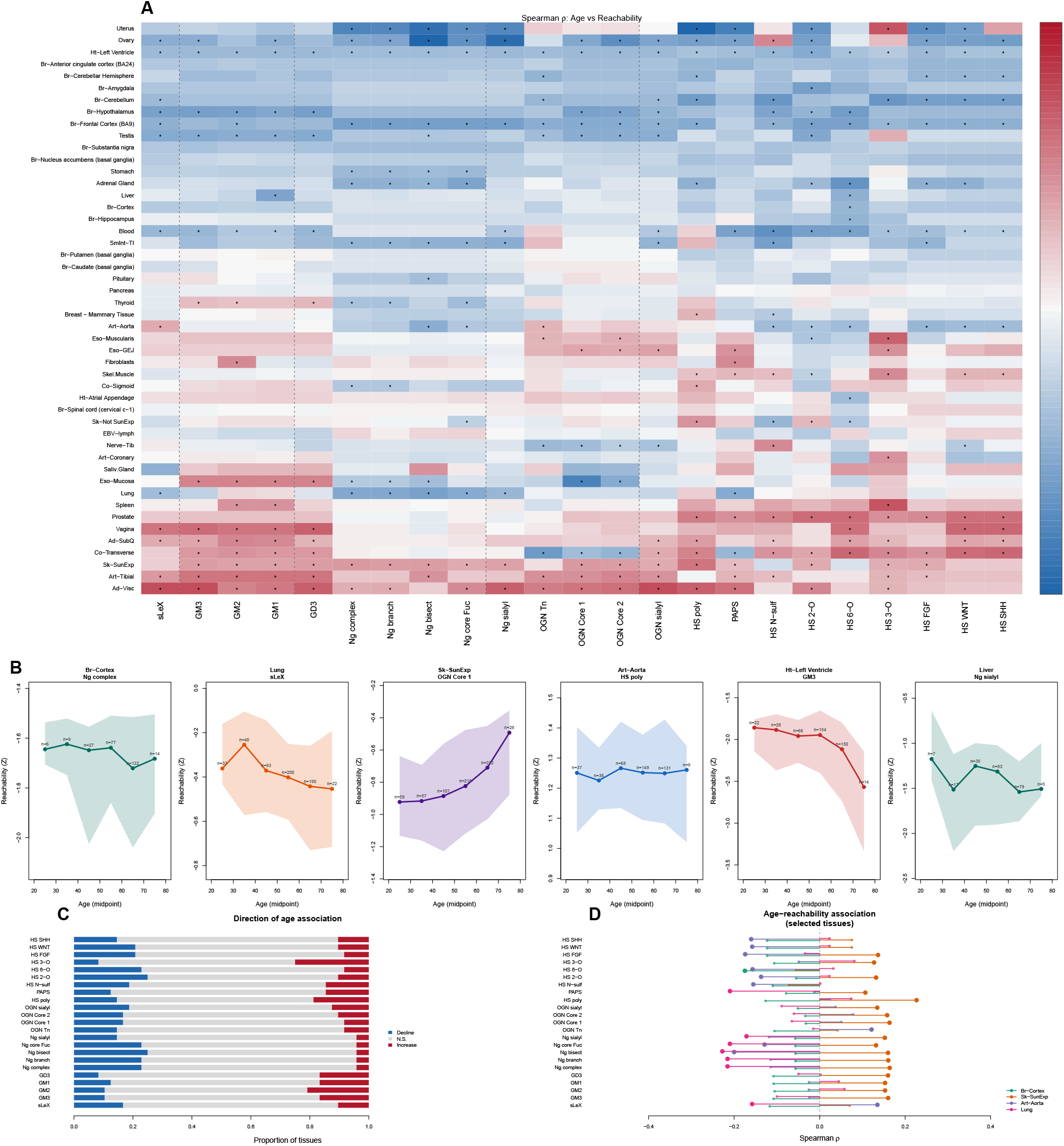
Age-related changes in glycan biosynthetic potential. **(A)** Summary of age-reachability Spearman correlations across 23 metrics and 48 evaluable tissues. Bar heights show the number of tissues with significant decline (blue, below) or increase (red, above) at FDR *<* 0.05. **(B)** Representative tissue-level age-reachability trajectories for selected metrics and tissues, showing the diversity of age-related patterns across organs.

N-glycan processing potential showed the most consistent age-related decline: Ng complex, Ng branch, and Ng bisect each showed significant decline in 13 of 48 evaluable tissues (FDR *<* 0.05), with median *ρ* ranging from −0.070 to −0.078 (Fig 4A). Ng core Fuc similarly declined in 13 tissues (median *ρ* = −0.067). These results are consistent with the known age-related reduction in N-glycan processing complexity observed in glycoproteomic studies.

Among the primary 14 metrics, HS 2-O sulfation showed the most tissues with significant age-related decline (12/48), followed by HS 6-O sulfation (11/48) and sLeX reachability (9/48). In contrast, HS 3-O sulfation was the only metric with more tissues showing significant increase (12/48) than decline (4/48), suggesting a distinct age-related regulatory program for this rare sulfation modification.

Ganglioside metrics showed mixed patterns: GM3 reach increased in 9 tissues but declined in 5, while GM1 and GD3 showed similar modest trends. The tissue-specificity of these age-related changes — varying in both direction and magnitude across organs — underscores the importance of sample-level analysis rather than aggregate summaries.

Results were essentially unchanged when controlling for sex via partial Spearman correlation (median Δ*ρ <* 0.01 across all tissue–metric pairs). RIN and ischemic time are not available as per-sample covariates in GTEx v8 open-access data.

We emphasize that the age-related effects reported here are small in magnitude (median |*ρ*| *<* 0.10 for most metrics) and should be considered exploratory. Age in GTEx is confounded with post-mortem interval, cause of death, and other donor-level variables that are incompletely controlled in the open-access data. These results generate testable hypotheses but do not establish causal age-related changes in glycan biosynthetic capacity.

### 3.7 Reachability is transcriptionally coherent with pathway-response genes beyond naive mean expression

We examined within-GTEx coherence between reachability and prespecified pathway-response transcript sets for WNT/HS, EGFR/GM3, and Selectin/sLeX across 50 tissues. Predictors and readouts come from the same RNA-seq matrix; accordingly, these analyses do not test glycan production, receptor engagement, or glycan-mediated causation, and tissue identity may confound them.

Tissue-level direction-oriented Spearman correlations were WNT *ρ* = 0.855, EGFR *ρ* = −0.718, and Selectin *ρ* = 0.564 (Fig 5A). For EGFR only, the response score was multiplied by −1 so that a positive coefficient would match the prespecified GM3-inhibitory direction; its observed negative coefficient therefore corresponds to a raw positive correlation and is evidence in the direction opposite that hypothesis. Corresponding naive all-pathway-gene means gave 0.719, −0.472, and 0.448. After controlling for the core-enzyme mean, direction-oriented partial correlations were 0.795, −0.743, and 0.508 (Fig 5B).

**Fig. 5:**
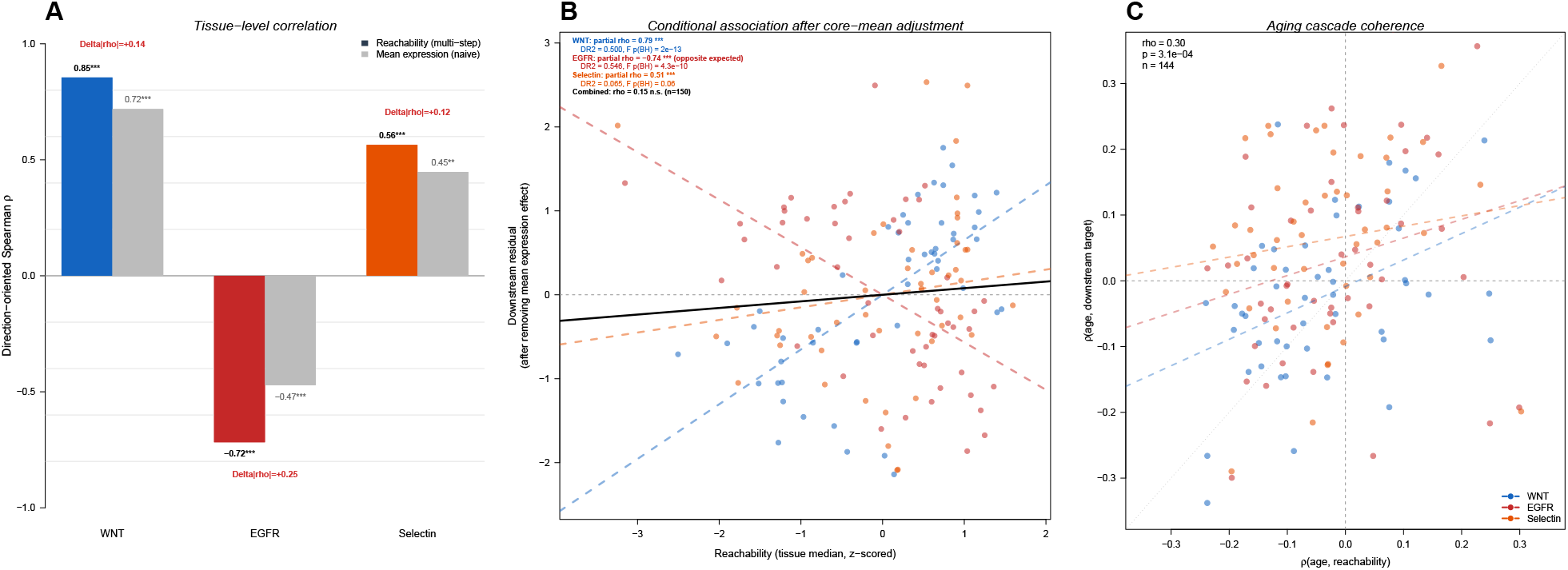
Reachability is transcriptionally coherent with pathway-response genes beyond naive mean expression. These are within-GTEx coherence analyses (predictors and readouts share an RNA-seq origin); they do not establish glycan-mediated causation (Section 3.7). **(A)** Tissue-level direction-oriented Spearman correlation (*ρ*) between each predictor and pathway-response gene expression for three independent cascades. Blue bars: reachability; grey bars: naive mean expression of all pathway genes. The EGFR response was multiplied by −1 so that positive would match the proposed GM3-inhibitory direction; its negative value is therefore opposite that direction. **(B)** Direction-oriented partial correlation after controlling for core mean expression; this is a conditional RNA-level association, not a unique causal contribution. **(C)** Aging coherence for 144 tissue-cascade pairs; both axes remain transcriptomic.

A variance-matched random-gene-set test retained WNT (*p* = 0.002) but not EGFR (*p* = 0.395) or Selectin (*p* = 0.783), consistent with generic tissue-expression covariance for the latter two. Other robustness checks are reported in Supplementary Methods and Supplementary File S2. These analyses describe the behavior of the constructed score on the same transcriptome and are not an external benchmark of biological accuracy.

Across 144 tissue–cascade combinations, age associations of reachability and response transcripts were also compared (Fig 5C). This is an additional within-transcriptome consistency analysis and provides no evidence that changes in glycans mediate the paired transcriptional trends.

### 3.8 Aggregation sensitivity on the within-GTEx coherence task

A central design choice is AND=min rather than averaging required steps. We compared the prespecified OR=mean/min score, an OR=max sensitivity, and two flat gene-expression averages (Fig 6). This analysis is explicitly post-hoc and internal to GTEx.

**Fig. 6:**
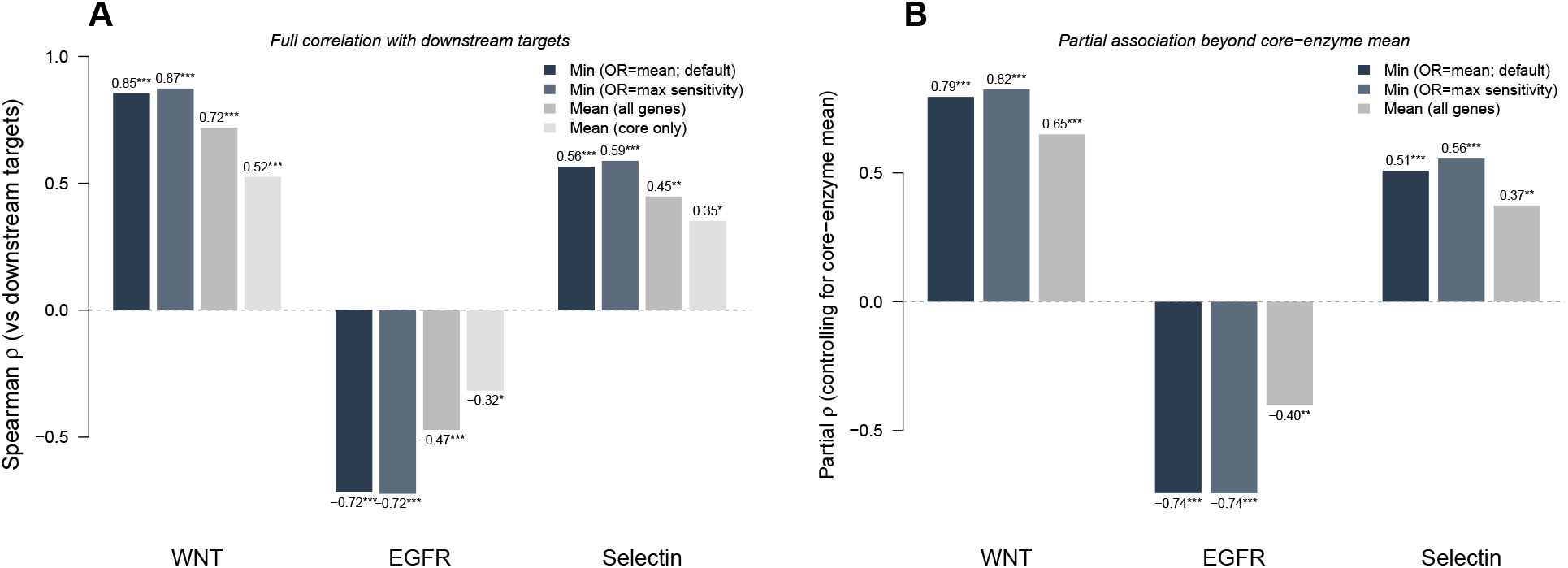
Aggregation sensitivity on the within-GTEx coherence task. **(A)** Direction-oriented Spearman correlations for the prespecified OR=mean/min score, OR=max sensitivity, all-gene mean, and core-only mean scores. **(B)** Direction-oriented partial correlations controlling for the core-only mean. EGFR response scores were multiplied by −1; negative EGFR values are opposite the prespecified inhibitory direction. These results compare internal RNA-level behavior and are not a glycan-accuracy benchmark.

The direction-oriented prespecified-score correlations were WNT 0.855, EGFR −0.718, and Selectin 0.564. OR=max gave 0.873, −0.722, and 0.588, while flat all-gene means gave 0.719, −0.472, and 0.448 (Fig 6A). These changes are small and internal to GTEx; they do not establish glycan-prediction accuracy. The generalized-quantile sweep is reported as a sensitivity analysis in S7 Fig.

### 3.9 External benchmark against knockout glycomics

We benchmarked three prespecified scorers on independent N-glycan mass-spectrometry profiles from wild-type and knockout HEK293 cells [4]: 41 N-glycans, six pathway-processing knockout strata, and an off-pathway B4GALNT3/4 negative control (Fig 7). The scorers were the minimum-based delta, a naive mean-aggregation delta, and GlycoMaple-style binary topology. None has fitted parameters, so reporting each fixed scorer over knockout strata is stratified external evaluation rather than cross-validation. AUROC, AUPRC, and Spearman correlation with observed | log_2_ FC| were computed per knockout.

**Fig. 7:**
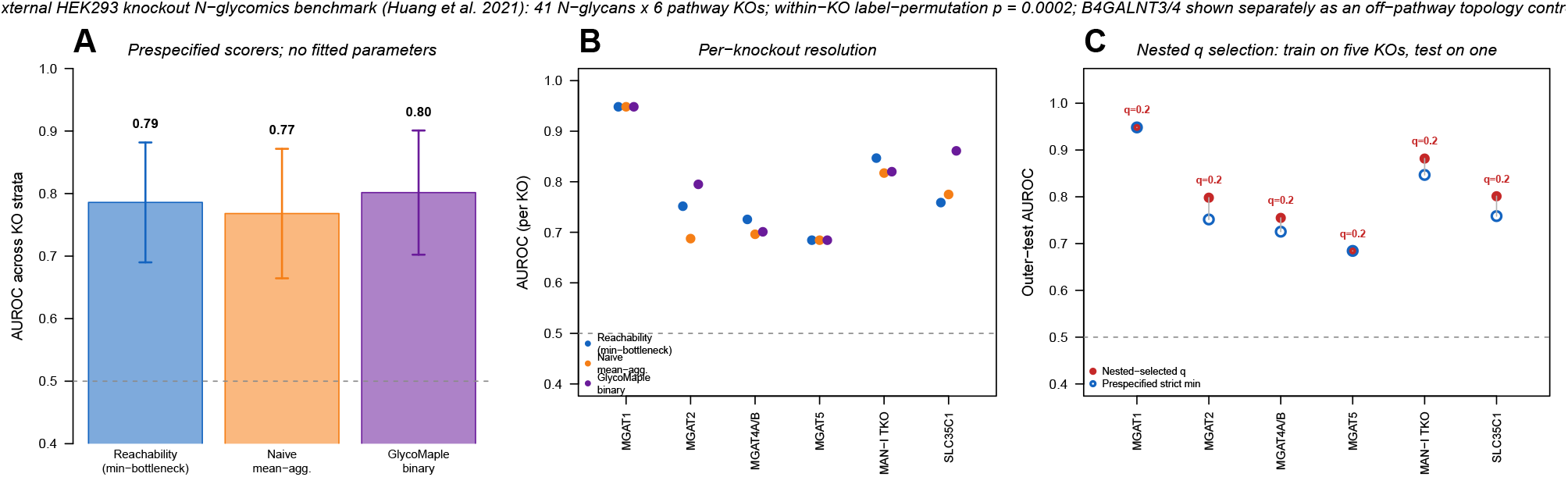
External benchmark against knockout glycomics. Three prespecified, parameter-free scorers were evaluated across six HEK293 knockout strata; B4GALNT3/4 was an off-pathway negative control. **(A)** Mean *±* s.d. AUROC across knockout strata. Minimum scoring did not outperform mean aggregation or binary topology. **(B)** Per-knockout AUROC, showing heterogeneity among perturbations. **(C)** For the only selected parameter, *q*, nested leave-one-knockout-out evaluation chose *q* on five knockouts and scored the sixth. Every fold selected *q* = 0.2 and outer-fold mean AUROC was 0.811, favoring a relaxed low quantile rather than validating the strict minimum.

Mean AUROC across the six strata was 0.786 for the minimum, 0.768 for the mean, and 0.802 for binary topology; mean AUPRC was 0.585, 0.582, and 0.591, respectively. Paired AUROC differences for minimum minus mean (0.018, bootstrap 95% CI [−0.003, 0.039], exact *p* = 0.25) and minimum minus binary (−0.016, [−0.054, 0.017], exact *p* = 0.625) did not support superiority of the minimum. Pooled minimum-delta AUROC was 0.820 and exceeded a label-permutation null stratified within knockout (*p* = 0.0002), indicating non-random information without relative superiority. The negative control had no predicted pathway pairs, although 3/37 glycans decreased and 5/37 changed by at least two-fold; median | log_2_ FC| was 0.24.

Aggregation quantile was the only selected parameter and was assessed with nested leave-one-knockout-out evaluation: *q* was chosen on five knockouts and scored once on the sixth. Every outer fold selected *q* = 0.2, yielding mean outer-fold AUROC 0.811. This favors a relaxed low quantile rather than validating the strict minimum. The external benchmark therefore supports a limited claim: the topology-aware score is informative above a null, but the available perturbation data do not show that the strict minimum is more accurate than the named baselines.

### 3.10 External tests against measured glycomes and essentiality: negative results and constraints

We performed a cell-line essentiality analysis and a tissue-level glycomic comparison. Neither provided positive support for tissue-level glycan-abundance ranking; these results constrain interpretation (see Discussion).

#### DepMap essentiality analysis (S4 Fig)

We tested whether glycosylation genes predicted as pathway bottlenecks show greater essentiality (more negative Chronos scores) in cancer cell lines from the DepMap/Project Achilles dataset 28]. Across the three cascades, only 2 of 23 tested bottleneck genes showed significantly different essentiality between bottleneck and non-bottleneck cell lines after BH correction: HS6ST1 (*p*_adj_ = 0.018) and HS6ST2 (*p*_adj_ = 0.012) in the WNT/HS cascade. The overall median Chronos scores for bottleneck vs. non-bottleneck cell lines were not meaningfully different for any cascade. This negative result is expected: glycan biosynthesis is generally non-essential for cancer cell proliferation in standard 2D culture conditions, as essentiality screens primarily detect growth/survival phenotypes rather than glycan-dependent functions such as cell-cell adhesion, immune recognition, and signaling modulation.

#### Cross-species tissue glycomics comparison (S5 Fig)

None of four comparisons with published mouse tissue glycomic profiles [42] was significant (all *p* ≥ 0.308, *n* = 15; Supplementary File S2). This provides no positive tissue-level abundance evidence.

## 4 Discussion

### 4.1 Relationship to existing methods

Glycan reachability analysis occupies a distinct methodological niche compared to existing computational glycobiology tools (Table 4). Unlike GlycoMaple [4], which answers “can this tissue produce glycan X?” (binary), our method answers “how does this tissuefs transcriptomic potential to produce glycan X compare to other tissues?” (quantitative, relative). Unlike glycoPATH [7], which predicts absolute glycan abundances via supervised machine learning requiring paired glycomic–transcriptomic training data, our method is unsupervised and requires only RNA-seq data. Unlike mechanistic models of glycosylation — ordinary-differential-equation and flux models [12, 13], the Markov-chain framework that learns isozyme specificities (e.g. GlycoMME) [14], and automated rule-based reaction-network construction [6, 15] — which require kinetic rate parameters or reaction rules rarely available at scale for most biological systems, our method relies on the biologically motivated assumption that the least-expressed pathway component limits overall pathway potential. Reachability thus occupies the low-parameter, cross-tissue-comparative extreme of this modelling continuum, complementary to (not competing with) the mechanistic pole.

**Table 3:** Kruskal–Wallis tests for tissue-level variation in reachability scores (14 primary metrics). All *p*-values are below machine precision (< 2.2 × 10^−308^).

| Metric | $\chi^2$ | df | $p$-value |
| --- | --- | --- | --- |
| sLeX reachability | 10,128 | 53 | $< 2.2 \times 10^{-308}$ |
| GM3 reach | 9,980 | 53 | $< 2.2 \times 10^{-308}$ |
| GM2 reach | 10,502 | 53 | $< 2.2 \times 10^{-308}$ |
| GM1 reach | 11,330 | 53 | $< 2.2 \times 10^{-308}$ |
| GD3 reach | 9,458 | 53 | $< 2.2 \times 10^{-308}$ |
| HS polymerization* | 14,649 | 53 | $< 2.2 \times 10^{-308}$ |
| PAPS | 13,113 | 53 | $< 2.2 \times 10^{-308}$ |
| HS N-sulfation | 10,991 | 53 | $< 2.2 \times 10^{-308}$ |
| HS 2-O sulfation | 10,407 | 53 | $< 2.2 \times 10^{-308}$ |
| HS 6-O sulfation | 11,561 | 53 | $< 2.2 \times 10^{-308}$ |
| HS 3-O sulfation | 13,125 | 53 | $< 2.2 \times 10^{-308}$ |
| HS FGF-like | 13,109 | 53 | $< 2.2 \times 10^{-308}$ |
| HS WNT-like | 12,673 | 53 | $< 2.2 \times 10^{-308}$ |
| HS SHH-like** | 12,398 | 53 | $< 2.2 \times 10^{-308}$ |

\*Uses AND (min) for EXT1/EXT2 heteromeric complex. \*\*Includes 3-O sulfation requirement (differs from WNT-like).

**Table 4:**
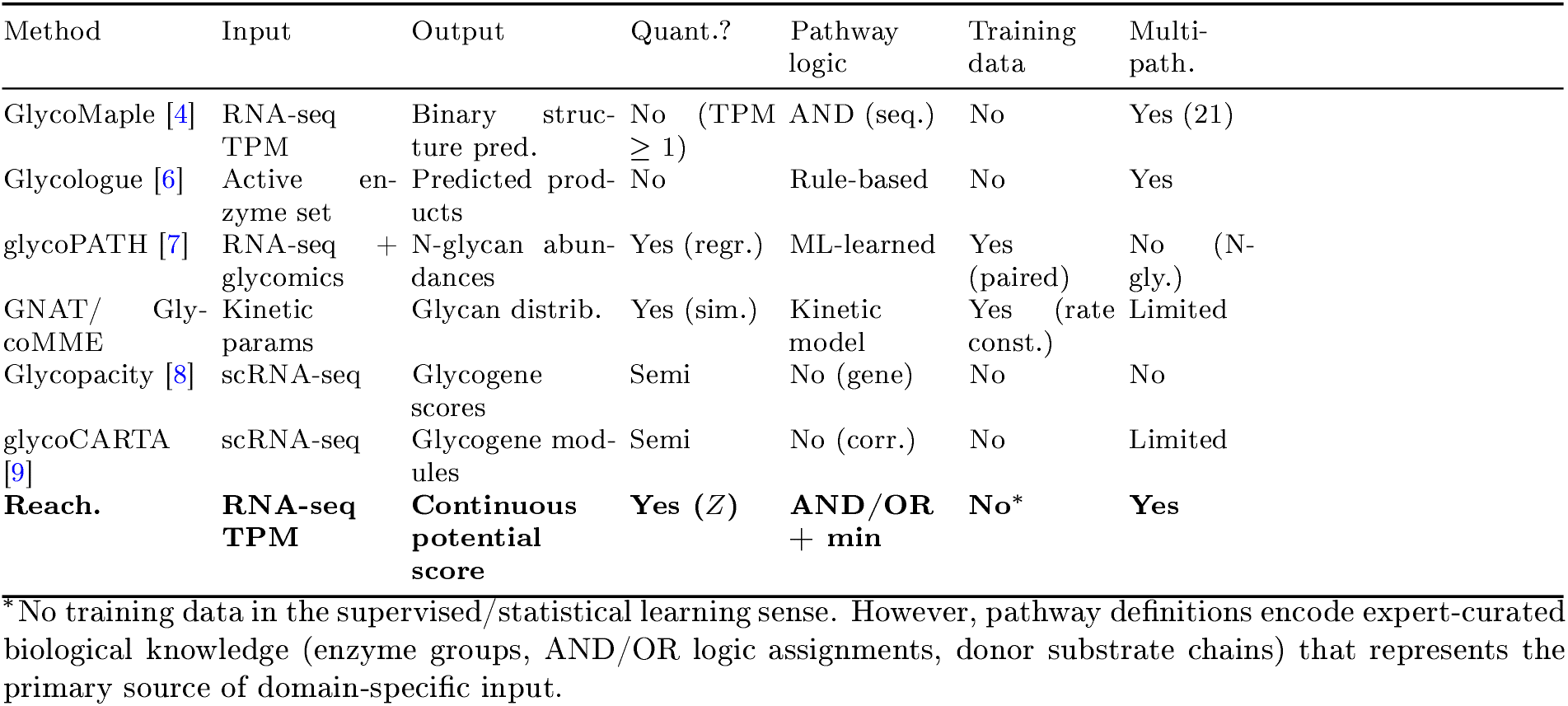
Comparison with existing computational glycobiology methods.

**Table 5:** Within-GTEx transcriptomic coherence statistics. These associations share an RNA-seq origin and are not biological validation. values are direction-oriented; negative EGFR coefficients are opposite the prespecified GM3-inhibitory direction. “Perm.” is the variance- matched random-gene-set test.

| Cascade | Metric | $\rho$ | 95% CI (tissue) | 95% CI (block) | Perm. $p$ |
| --- | --- | --- | --- | --- | --- |
| WNT | Full $\rho$ (reach) | 0.855 | [0.750, 0.908] | [0.852, 0.873] | 0.002 |
| WNT | Full $\rho$ (mean) | 0.719 | [0.511, 0.857] | — | — |
| WNT | Partial $\rho$ | 0.795 | [0.616, 0.879] | — | — |
| EGFR | Full $\rho$ (reach) | -0.718 | [-0.826, -0.549] | [-0.742, -0.700] | 0.395 |
| EGFR | Full $\rho$ (mean) | -0.472 | [-0.714, -0.182] | — | — |
| EGFR | Partial $\rho$ | -0.743 | [-0.853, -0.543] | — | — |
| Selectin | Full $\rho$ (reach) | 0.564 | [0.311, 0.740] | [0.536, 0.581] | 0.783 |
| Selectin | Full $\rho$ (mean) | 0.448 | [0.168, 0.666] | — | — |
| Selectin | Partial $\rho$ | 0.508 | [0.173, 0.721] | — | — |

The bottleneck (min-aggregation) principle is a heuristic motivated by linear-regime enzyme kinetics. Metabolic control analysis instead shows that flux control can be distributed among multiple enzymes [35, 36]; expression is also not enzyme activity. Our method therefore identifies an expression argmin, not a biochemical rate-limiting step. The stability analysis shows whether that argmin recurs among donors. Nested external selection favored *q* = 0.2, while the strict minimum did not outperform the mean or binary baselines. The evidence supports interpretability and non-random signal, not optimality of the strict minimum.

While conceptually related to constraint-based metabolic modeling approaches such as FBA [24], our method differs fundamentally in scope. FBA operates on genome-scale stoichiometric matrices with mass conservation constraints and linear programming. Expression-constrained approaches such as iMAT [25] and GIMME [26] share our goal of integrating transcriptomic data with pathway knowledge, but require genome-scale metabolic models. The trade-off is explicit: our method sacrifices mechanistic detail for simplicity, interpretability, and broad applicability to any pathway definable with AND/OR logic.

The aggregation comparison (Fig 6) shows that min-aggregation is more correlated with the chosen response transcripts on this within-GTEx task. Because the target sets, predictors, and scoring data share a transcriptomic source, this is not evidence that the minimum predicts glycans better. The independent knockout-glycomics benchmark (Section 3.9) did not establish such superiority either.

### 4.2 Biological interpretation of discordant cases

The case studies below illustrate two systematic failure modes of bulk tissue reachability analysis cell-type heterogeneity masking minority populations, and cell-type composition differences mimicking biosynthetic differences — that define the interpretive boundaries of our approach.

The pancreas case (Fig 2C) exemplifies where quantitative assessment provides additional resolution. While a binary approach classifies pancreas as capable of sLeX synthesis (96% of samples have all enzymes detectable), the quantitative reachability score (*Z* = −1.86) indicates uniformly low expression across all pathway steps. The exocrine pancreas, which dominates the bulk tissue transcriptome, primarily produces digestive enzymes and maintains glycosylation machinery at basal levels. We note, however, that pancreatic ductal cells — a minority population — are known to express sLeX and related antigens at higher levels, and this cell-type heterogeneity limits the interpretation of bulk tissue predictions for the pancreas.

The brain ganglioside case (Fig 2D) is a failure case for abundance interpretation. Brain is the most ganglioside-rich organ [37], whereas bulk-brain GM3 reachability is low and its argmin identity is unstable. Cell composition is a plausible explanation, but it was not tested here and should not be used to dismiss the discrepancy. The result is genuine counter-evidence to using bulk-tissue reachability as a general abundance proxy and motivates cell-resolved evaluation.

### 4.3 External comparisons and limitations

We weight the external comparisons by their relevance. The DepMap null is a scope boundary because proliferation screens may miss glycan phenotypes. The mouse tissue-glycome comparison is more consequential and was null. Together with the brain discrepancy, it lowers confidence that reachability is a useful proxy for realized abundance across bulk tissues. The HEK293 perturbation analysis shows above-null topology-related information, not validation of cross-tissue abundance ranks.

The GTEx coherence analysis is circular as a biological evaluation: predictors and response transcripts are derived from the same RNA-seq. It cannot establish glycan production or mediation, and both tissue-level and within-tissue correlations remain vulnerable to cell composition and other co-regulation. Donor-block bootstrap intervals and within-tissue checks describe precision and robustness of the RNA-level association, not causal validity (Supplementary File S2). Shared donors can also make pairwise tissue-test *p*-values anti-conservative, so effect sizes and within-metric FDR are emphasized.

Gene expression does not measure enzymatic activity, protein stability, localization, substrate availability, or product turnover. The deterministic minimum also lacks an uncertainty term for annotation error, redundancy, compensation, or enzyme promiscuity. FUT8 is the canonical mammalian core *α*1,6-fucosyltransferase, yet a platelet-specific deletion study reported residual core-fucosylated signal in the perturbed context [43]; whether this reflects incomplete deletion, analytical assignment, residual protein, or unmodeled activity, a hard zero is not a safe quantitative prediction. Both the internal sweep and nested external selection favored a relaxed low quantile over the strict minimum. We therefore interpret the strict minimum as an ordinal expression heuristic, never a hard ceiling or zero-production rule. Glycosidase trimming, Golgi organization, post-mortem effects [27], and bulk-tissue heterogeneity remain unmodeled. A probabilistic per-step extension that propagates expression and annotation uncertainty is therefore a priority for future work.

The current implementation covers five glycan families; expansion requires additional curation. Within the HS pathway, the HS3ST isoforms (HS3ST1-6) are modeled as a single OR group despite having distinct substrate specificities [33], and the 6-O endosulfatases SULF1/SULF2, which modulate WNT ligand binding [30], are omitted. The model also omits inter-pathway competition for shared nucleotide sugar donors (CMP-Sia, UDP-Gal, GDP-Fuc), two donor pathways (UDP-GlcNAc, UDP-GlcA), the HS linker tetrasaccharide (XYLT1/2, B4GALT7, B3GALT6, B3GAT3), and acceptor protein availability (e.g., PSGL-1 for sLeX, syndecans/glypicans for HS). Finally, Z-scores are relative to the GTEx population, making scores dataset-dependent, and min-aggregation produces systematically negative scores for pathways with many steps, precluding direct cross-pathway comparison (see Section 3.2 for details).

### 4.4 Perspectives

The score nominates, rather than predicts, cases for direct measurement. For example, pancreas combines near-universal binary detectability with low sLeX reachability, whereas the brain result conflicts with known ganglioside abundance. Matched glycomic–transcriptomic measurements in the same specimens and cell types are needed to determine when either ranking is useful. Age associations likewise generate hypotheses and should not be interpreted as changes in glycan abundance without direct measurement.

The argmin can guide a falsifiable perturbation experiment: upregulating the nominated step and a matched non-argmin step would test whether the former changes product more. The HEK293 benchmark does not establish this claim because the minimum did not outperform the baselines. Prospective CRISPRa experiments with glycomic readouts are required.

Beyond the evaluations reported here, the framework enables several classes of biological inquiry. Applied to disease transcriptomes, reachability analysis could identify pathways whose biosynthetic potential shifts between healthy and pathological states — for instance, whether tumor-associated glycan changes (e.g., increased sLeX, altered branching) reflect transcriptomic reprogramming of specific bottleneck steps. Application to single-cell atlases would resolve the bulk tissue heterogeneity limitation and reveal cell-type-specific glycan biosynthetic programs, addressing cases such as the brain ganglioside paradox described above. Integration with spatial transcriptomics could further map glycan biosynthetic potential onto tissue architecture, linking reachability gradients to functional microenvironments. An R package implementing all methods described here is available at https://github.com/matsui-lab/glycoreach.

## Supporting information

Supplementary File S1: Pathway definitions

Supplementary File S2: Benchmark and sensitivity analyses

## Supplementary Methods

### Isozyme grouping and OR aggregation sensitivity

#### Isozyme (OR) aggregation and near-silent members

ST6GAL2 had mean tissue-median TPM 3.3 and was below 1 TPM in 29/54 tissues, versus 22.1 TPM for ST6GAL1. We used OR=mean as the primary definition and evaluated sensitivity to OR=max and ST6GAL1-only aggregation by recomputing Ng_sia under both alternatives. Tissue rankings were highly concordant (Spearman *ρ* = 0.993 and 0.960, respectively), although OR=max changed individual tissues by as much as 0.580 *Z*. The conclusions are therefore not driven by ST6GAL2 dilution, but the affected tissues are flagged in Supplementary File S2. Required complexes use AND=min.

#### Multiple testing correction strategy

BH correction was applied within each prespecified analysis family rather than across scientifically different tasks. The largest family comprised 28,175 pairwise Wilcoxon tests (1,225 tissue pairs *×* 23 metrics), corrected separately within each metric. Coherence panels and age analyses were corrected within their stated panels or tissue strata; empirical permutation *p*-values were not BH-adjusted. Sensitivity grids were treated descriptively and were not used as confirmatory hypothesis families. Supplementary File S2 enumerates the analysis families and corrections.

### Transcriptomic-coherence sensitivity analyses

#### Variance-matched permutation

For each cascade, 1,000 random gene sets of the same size as the response set were sampled with weights favoring genes with similar cross-tissue variance. Empirical two-sided *p*-values were 0.002, 0.395, and 0.783 for WNT, EGFR, and Selectin. This controls one feature of generic tissue covariance but does not create an external test.

#### Bootstrap confidence intervals

95% bootstrap confidence intervals (2,000 iterations, tissue-level resampling) were computed for all Spearman correlations and partial correlations reported in Panels A and B.

#### All-pathway-expression control

In addition to the prespecified core-enzyme control shown in Fig 5B, we controlled for the mean Z-score of every gene in each pathway definition. Partial correlations were 0.680, −0.641, and 0.387 for WNT, EGFR, and Selectin, respectively (nominal *p* = 7.9 *×* 10^−8^, 7.1 *×* 10^−7^, and 0.0060). This sensitivity remains entirely within the same RNA-seq data and is not an orthogonal validation.

#### Nonlinearity assessment

For the Selectin cascade, where the Spearman partial correlation was significant but the linear F-test was not, we fitted a generalized additive model (GAM; `mgcv` R package) with a smooth term for reachability and compared against the linear model using an F-test.

#### Pseudocount sensitivity analysis

We tested pseudocounts 0.1, 0.5, 1, and 5 (S9 Fig). This assesses numerical sensitivity of the coherence correlations only.

#### Normalization sensitivity analysis

Default log(1 + TPM) Z-scores gave direction-oriented correlations 0.855, −0.718, and 0.564; rank-based inverse-normal transformation gave 0.874, −0.718, and 0.553. Quantile normalization gave 0.535, 0.032, and −0.353, showing that the analysis is not invariant to normalization (S9 Fig).

### Aggregation function comparison

We compared four scoring constructions on the within-GTEx coherence task; this was not treated as glycan prediction accuracy:

- **Min-aggregation (OR=mean):** primary specification.
- **Min-aggregation (OR=max):** sensitivity specification.
- **Mean (all genes):** Flat mean of Z-scores across all pathway genes (both catalytic enzymes and substrate supply genes).
- **Mean (core only):** Mean of Z-scores for core catalytic enzymes only (ctrl_genes).

For each method-cascade combination, tissue-level Spearman correlation was computed (*n* = 50); partial correlations controlled for the core-only mean. These are conditional transcriptomic associations, not unique biological prediction.

### External knockout-glycomics benchmark

We used published N-glycan mass-spectrometry profiles of wild-type and knockout HEK293 cells [4]. The analysis included 41 N-glycan compositions and six processing knockouts — MGAT1, MGAT2, MGAT4A/B, MGAT5, the MAN1A1/A2/B1 mannosidase-I triple knockout, and SLC35C1 — with off-pathway B4GALNT3/4 retained as a negative control. Detectable glycans (WT abundance *>* 0.1) were labeled as decreased when observed log_2_ fold-change was *<* −1 (and as changed in either direction when | log_2_ FC| *>* 1); AUROC/AUPRC use the decrease label, whereas Spearman correlation uses the continuous | log_2_ FC|.

We compared three scorers that differ only in how a glycanfs required biosynthetic steps are aggregated: (i) *reachability* (min bottleneck), the drop in the minimum step level when the knocked-out genefs expression is set to zero; (ii) *naive ean-aggregation*, the analogous drop in the mean step level; and (iii) *GlycoMaple-style binary topology*, an indicator of whether the glycan requires the knocked-out gene (WT TPM ≥ 1). Each glycanfs required steps were assigned from its monosaccharide composition, with the antenna GlcNAc count parsed as the count immediately preceding the trimannosyl core. Genes absent from the HEK293 expression panel were treated as missing (excluded from aggregation) rather than as zero expression, so that a single unmeasured gene could not spuriously collapse the minimum; genes measured as zero were retained.

The three fixed scorers have no fitted parameter. They were evaluated separately in each of six knockout strata using AUROC, AUPRC, and Spearman correlation with observed | log_2_ FC|; foldwise AUROC differences were compared by exact sign permutation and bootstrap intervals. A 5,000-iteration label-permutation null shuffled labels within knockout. Quantile *q* was the only selected parameter and was assessed by nested leave-one-knockout-out evaluation: each outer knockout was scored once after *q* was chosen using the other five. This distinction prevents fixed-scorer stratification from being mislabeled as cross-validation.

### Additional sensitivity analyses

#### Leave-one-tissue-group-out analysis

To assess whether correlated tissue groups (e.g., multiple brain regions) drive the coherence results, we systematically removed all tissues belonging to each of three groups — brain (*n* = 13), gastrointestinal (*n* = 7), and reproductive (*n* = 5) — and recomputed tissue-level Spearman correlations for all three cascades.

#### Block bootstrap

To account for non-independence among related tissues, we performed block bootstrap with 2,000 iterations, resampling tissue groups (brain, artery, skin, esophagus, adipose, heart, colon, kidney, and remaining tissues as individual blocks) with replacement, and computed 95% confidence intervals.

#### Glycosidase expression survey

To assess whether glycan-degrading enzymes confound reachability predictions, we surveyed the expression of 12 glycosidases — categorized by subcellular localization (lysosomal: NEU1, MAN2B1/2, FUCA1, HEXA/B, GBA, GALC; plasma membrane: NEU3, NEU4; cytoplasmic: NEU2; ER/Golgi: FUCA2) — and computed Spearman correlations with corresponding reachability metrics across 50 tissues (S8 Fig).

#### Donor substrate demand analysis

To assess the potential impact of inter-pathway competition for shared nucleotide sugar donors, we computed aggregate demand indices for CMP-Sia, UDP-Gal, and GDP-Fuc per tissue by summing the tissue-median Z-scores of all consuming enzymes across all five pathway families (S1 Table).

#### Generalized aggregation analysis

AND aggregation was swept over 21 quantiles from minimum to maximum while OR=mean was held fixed. The same GTEx response correlations were used only to describe sensitivity; the sweep was not used for model selection (S7 Fig).

#### Donor-level block bootstrap

To assess the impact of donor-level non-independence in GTEx (948 donors contributing to multiple tissues), we performed donor-level block bootstrap: 2,000 iterations resampling donors (not tissues) with replacement, reconstructing tissue medians from each resampled donor set, and computing Spearman correlations (S2 Table).

#### Tissue-subtype collapsing

To estimate the effective sample size after accounting for non-independent tissue subtypes, we collapsed the 50 tissues to approximately 30 organ-level representatives (e.g., all 13 brain regions to a single “Brain”, 3 arteries to “Artery”) and recomputed tissue-level correlations at this reduced sample size (S2 Table).

#### Within-tissue sample-level correlations

To test whether the tissue-level associations hold at the individual sample level (addressing the ecological fallacy), we computed within-tissue Spearman correlations between per-sample reachability and per-sample downstream target expression for each of the 50 tissues and each cascade (S2 Table).

#### Bottleneck stability analysis

All 19 multi-step metrics were evaluated in each of 50 sufficiently sampled tissues. We computed modal-argmin frequency, normalized entropy, the median gap between the smallest and second-smallest step, 1,000 random donor-half splits, and 1,000 donor bootstrap resamples. Overall median modal frequency was 0.608; mean half-split agreement was 0.917 and mean bootstrap recovery 0.958. Four one-step endpoints were excluded because argmin stability is not applicable. Full tissue-metric results are in Supplementary File S2.

#### Steiger’s test for correlation differences

To formally test whether reachability is significantly more strongly correlated than mean expression, we applied the Steiger (1980) test for comparing two dependent correlations sharing a common variable (the pathway-response readout) (S2 Table).

#### Restricted response-gene sensitivity

We repeated the coherence analysis with WNT: AXIN2/LEF1; EGFR: AREG/EREG/EGR1; and Selectin: SELPLG/ITGB2/ICAM1 (Supplementary File S2).

## Data and Code Availability

GTEx v8 gene-expression data and annotations are publicly available from the GTEx Portal (https://gtexportal.org/); the exact public URLs and checksums are encoded in `reproducibility/fetch_large_data.sh`. HEK293 KO glycomics were obtained from 4]; DepMap data are identified by release in the reproduction manifest. Analysis scripts, environment specifications, derived per-sample reachability, statistical outputs, and figure source data are available at https://github.com/matsui-lab/glycoreach. The immutable reproducibility package (version `manuscript-r1-v1.0.0`) is archived at Zenodo (https://doi.org/10.5281/zenodo.21428810). Supplementary File S1 contains pathway definitions and Supplementary File S2 contains the complete benchmark and sensitivity outputs, including data behind reported means, medians, intervals, and figures. `reproducibility/README_REPRODUCE.md` describes both a no-large-download figure rebuild and a raw-data rebuild.

## Acknowledgments

The author thanks the GTEx donors and consortium and the investigators who made the glycomic datasets publicly available.

## Supporting Information

**S1 Fig**. Dimensionality reduction of glycogene expression.

**S2 Fig**. Full glycogene expression landscape.

**S3 Fig**. Within-GTEx correlations with receptor and pathway-response transcripts.

**S4 Fig**. DepMap essentiality analysis (negative result).

**S5 Fig**. Cross-species (mouse) tissue glycomics comparison (negative result).

**S6 Fig**. Reachability versus naive mean expression.

**S7 Fig**. Generalized aggregation analysis (min-to-max quantile sweep).

**S8 Fig**. Glycosidase expression survey.

**S9 Fig**. Normalization and pseudocount sensitivity.

**S10 Fig**. Tissue median reachability landscape.

**S11 Fig**. Per-knockout descriptive detail for the knockout-glycomics benchmark.

**S1 Table**. Donor substrate demand indices.

**S2 Table**. Summary of within-GTEx coherence sensitivities.

**S3 Table**. GTEx v8 tissue sample sizes.

**S4 Table**. Master list of all 23 reachability metrics.

**S5 Table**. Tissue name abbreviations used in figures.

**S1 File**. Machine-readable glycan pathway definitions (XLSX).

**S2 File**. Complete benchmark, bottleneck-stability, isozyme, tissue-glycome, and multipletesting outputs (XLSX).

## References

[1] Neelamegham S, Aoki-Kinoshita KF, Bolton E, Frank M, Lisacek F, Lütteke T, et al. Updates to the Symbol Nomenclature for Glycans guidelines. Glycobiology. 2019;29(9):620–624.

[2] Groth T, Diehl AD, Gunawan R, Neelamegham S. GlycoEnzOnto: a GlycoEnzyme pathway and molecular function ontology. Bioinformatics. 2022;38(24):5413–5420.

[3] arki A. Biological roles of glycans. Glycobiology. 2017;27(1):3–49.

[4] Huang YF, Aoki K, Akase S, Ishihara M, Liu YS, Yang G, et al. Global mapping of glycosylation pathways in human-derived cells. Dev Cell. 2021;56(8):1195–1209.

[5] Kong WZ, Fujita M. GlycoMaple: recent updates and applications in visualization and analysis of glycosylation pathways. Anal Bioanal Che . 2025;417:885–894.

[6] McDonald AG, Tipton KF, Davey GP. A knowledge-based system for display and prediction of O-glycosylation network behaviour in response to enzyme activities. PLoS Comput Biol. 2016;12(4):e1004844.

[7] Alvarez MRS, Holmes XA, Oloumi A, Grijaldo-Alvarez SJ, Schindler R, Zhou Q, et al. Integration of RNAseq transcriptomics and N-glycomics reveal biosynthetic pathways and predict structure-specific N-glycan expression. Che Sci. 2025;16:7155–7172.

[8] Dworkin LA, Clausen H, Joshi HJ. Applying transcriptomics to study glycosylation at the cell type level. iScience. 2022;25(6):104419.

[9] Chrysinas P, Venkatesan S, Ang I, Ghosh Chen C, Neelamegham S, et al. Cell- and tissue-specific glycosylation pathways informed by single-cell transcriptomics. NAR Ge-no ics Bioinf. 2024;6(4):qae169.

[10] J GTEx Consortium. The GTEx Consortium atlas of genetic regulatory effects across human tissues. Science. 2020;369(6509):1318–1330.

[11] Nairn A , York WS, Harris K, Hall EM, Pierce JM, Moremen KW. Regulation of glycan structures in animal tissues: transcriptome analysis of glycosyltransferase expression. J Biol Che . 2008;283(25):17298–17313.

[12] Uma a P, Bailey JE. A mathematical model of N-linked glycoform biosynthesis. Biotechnol Bioeng. 1997;55(6):890–908.

[13] Krambeck FJ, Bennun S , Narang S, Barnard J, Goldman JM, Betenbaugh MJ. A mathematical model to derive N-glycan structures and cell surface glycoprotein activities from mass spectrometric data. Glycobiology. 2009;19(11):1163–1175.

[14] Spahn PN, Hansen AH, Hansen HG, Arnsdorf J, Kildegaard HF, Lewis NE. A Markov chain model for N-linked protein glycosylation towards a low-parameter tool for model-driven glycoengineering. Metab Eng. 2016;33:52–66.

[15] Liu G, Neelamegham S. A computational framework for the automated construction of glycosylation reaction networks. PLoS One. 2014;9(6):e100939.

[16] Janda CY, Waghray D, Levin AM, Thomas C, Garcia KC. Structural basis of Wnt recognition by Frizzled. Science. 2012;337(6090):59–64.

[17] Nusse R, Clevers H. Wnt/β-catenin signaling, disease, and emerging therapeutic modalities. Cell. 2017;169(6):985–999.

[18] Kawashima N, Yoon SJ, Itoh K, Nakayama KI. Tyrosine kinase activity of epidermal growth factor receptor is regulated by GM3 binding through carbohydrate to carbohydrate interactions. J Biol Che . 2009;284(10):6147–6155.

[19] Yoon SJ, Nakayama KI, Hikita T, Handa K, Hakomori SI. Epidermal growth factor receptor tyrosine kinase is modulated by GM3 interaction with N-linked GlcNAc termini of the receptor. Proc Natl Acad Sci USA. 2006;103(49):18987–18991.

[20] Pratilas CA, Taylor BS, Y. , iale A, Sander C, Solit DB, et al. 600EBRAF is associated with disabled feedback inhibition of RAF-MEK signaling and elevated transcriptional output of the pathway. Proc Natl Acad Sci USA. 2009;106(11):4519–4524.

[21] Ley K, Laudanna C, Cybulsky MI, Nourshargh S. Getting to the site of inflammation: the leukocyte adhesion cascade updated. Nat Rev Immunol. 2007;7(9):678–689.

[22] Kim S. ppcor: an R package for a fast calculation to semi-partial correlation coefficients. Co un Stat Si ul Co put. 2015;44(4):1062–1078.

[23] Gagiannis D, Gossrau R, Reutter W, Zimmermann-Kordmann M, Horstkorte R. Engineering the sialic acid in organs of mice using N-propanoylmannosamine. Biochi Biophys Acta. 2007;1770(2):297–306.

[24] Orth JD, Thiele I, Palsson BO. What is flux balance analysis? Nat Biotechnol. 2010;28(3):245–248.

[25] Shlomi T, Cabili MN, Herrgiård MJ, Palsson BO, Ruppin E. Network-based prediction of human tissue-specific metabolism. Nat Biotechnol. 2008;26(9):1003–1010.

[26] Becker SA, Palsson BO. Context-specific metabolic networks are consistent with experiments. PLoS Comput Biol. 2008;4(5):e1000082.

[27] Ferreira PG, Mu oz-Aguirre M, Reverter F, Sa Godinho CP, Sousa A, Amadoz A, et al. The effects of death and post-mortem cold ischemia on human tissue transcriptomes. Nat Co un. 2018;9(1):490.

[28] Dempster JM, Boyle I, Vazquez F, Root DE, Boehm JS, Hahn WC, et al. Chronos: a cell population dynamics model of CRISPR experiments that improves inference of gene fitness effects. Geno e Biol. 2021;22(1):343.

[29] Rapraeger AC, Krufka A, Olwin BB. Requirement of heparan sulfate for bFGF-mediated fibroblast growth and myoblast differentiation. Science. 1991;252(5013):1705–1708.

[30] Ai X, Do AT, Lozynska O, Kusche-Gullberg M, Lindahl U, Emerson CP Jr. Sulf1 remodels the 6-O sulfation states of cell surface heparan sulfate proteoglycans to promote Wnt signaling. J Cell Biol. 2003;162(2):341–351.

[31] McCormick C, Duncan G, Goutsos KT, Tufaro F. The putative tumor suppressors EXT1 and EXT2 form a stable complex that accumulates in the Golgi apparatus and catalyzes the synthesis of heparan sulfate. Proc Natl Acad Sci USA. 2000;97(2):668–673.

[32] Openo KK, Schulz JM, Vargas CA, Orton CS, Epstein MP, Schnur RE, et al. Epimerase-deficiency galactosemia is not a binary condition. A J Hu Genet. 2006;78(1):89–102.

[33] Shworak NW, Liu J, Petros LM, Zhang L, Kobayashi M, Copeland NG, et al. Multiple isoforms of heparan sulfate D-glucosaminyl 3-O-sulfotransferase. J Biol Che . 1999;274(9):5170–5184.

[34] Romano J, Kromrey JD, Coraggio J, Skowronek J. Appropriate statistics for ordinal level data: Should we really be using t-test and Cohenfs d for evaluating group differences on the NSSE and similar surveys? In: Annual Meeting of the Florida Association of Institutional Research; 2006:1–33.

[35] Kacser H, Burns JA. The control of flux. Sy p Soc Exp Biol. 1973;27:65–104.

[36] Heinrich R, Rapoport TA. A linear steady-state treatment of enzymatic chains. General properties, control and effector strength. Eur J Biochem. 1974;42(1):89–95.

[37] Schnaar RL. Gangliosides of the vertebrate nervous system. J Mol Biol. 2016;428(16):3325–3336.

[38] Ballehaninna UK, Chamberlain RS. The clinical utility of serum CA 19-9 in the diagnosis, prognosis and management of pancreatic adenocarcinoma: an evidence based appraisal. J Gastrointest Oncol. 2012;3(2):105–119.

[39] Cliff N. Dominance statistics: Ordinal analyses to answer ordinal questions. Psychol Bull. 1993;114(3):494–509.

[40] Efron B, Tibshirani RJ. An Introduction to the Bootstrap. New York: Chapman Hall; 1993.

[41] Phipson B, Smyth GK. Permutation P-values should never be zero: calculating exact P-values when permutations are randomly drawn. Stat Appl Genet Mol Biol. 2010;9(1):Article 39.

[42] Otaki M, Hirane N, Natsume-Kitatani Y, Nogami Itoh M, Shindo M, Kurebayashi Y, et al. Mouse tissue glycome atlas 2022 highlights inter-organ variation in major N-glycan profiles. Sci Rep. 2022;12(1):17804.

[43] Yang RB, Tu CF, Chen YT, Chau TH, Lin SW, Tsai CD, et al. FUT8-dependent core fucosylation: essential for platelet function and a target in thrombosis. Arterioscler Thro b Vasc Biol. 2026. doi:10.1161/ATBAHA.126.324757.

