## Supporting Information for "Glycan Reachability Analysis: A Bottleneck-Aware Framework for Inferring Tissue-Specific Glycan Biosynthetic Potential from Transcriptomics"

Yusuke Matsui

#### Supporting Information

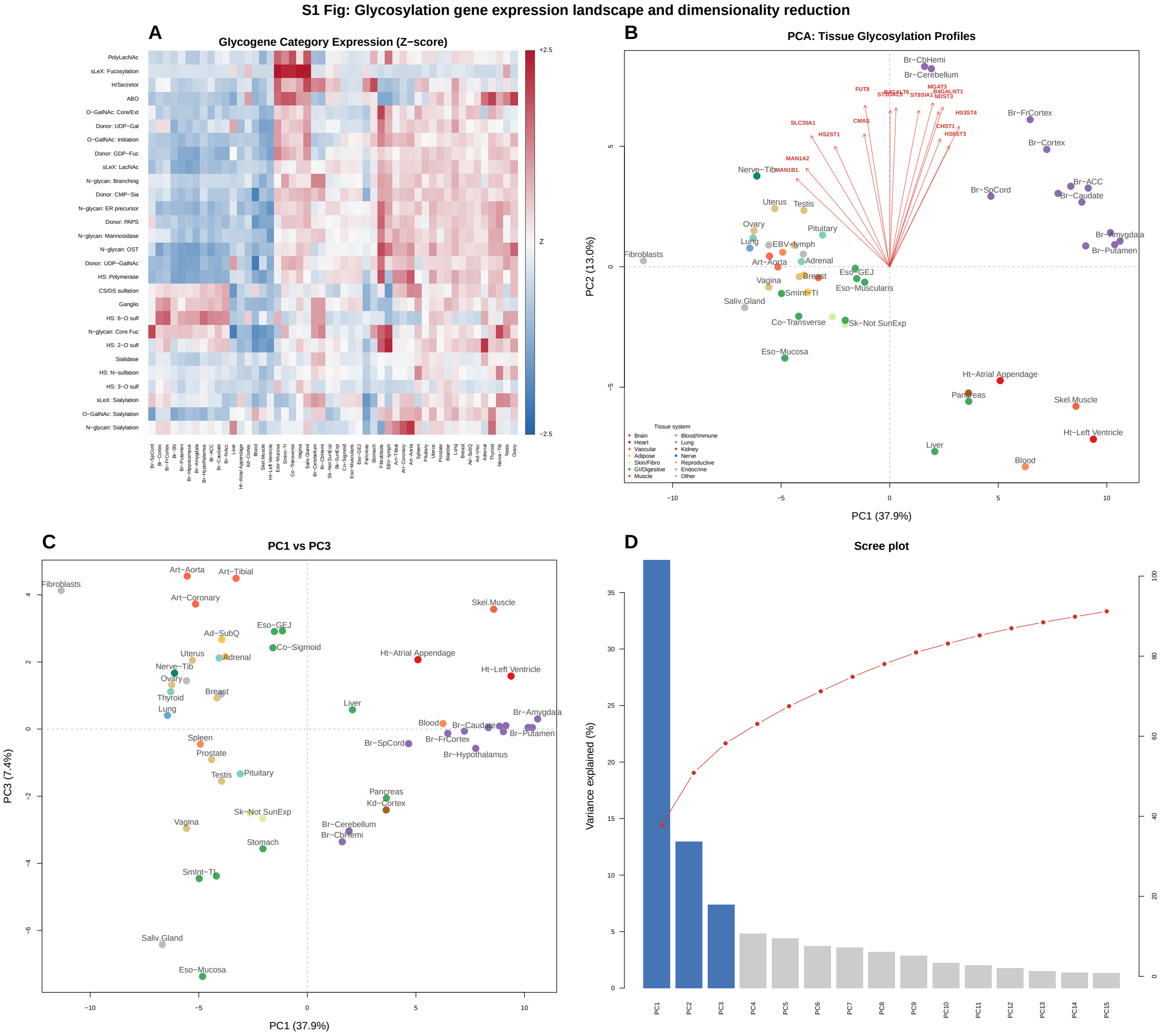


**S1 Fig. Dimensionality reduction of glycogene expression. (A)** Heatmap of median Z-scored glycogene expression across 50 GTEx tissues (rows: 25 gene categories; columns: tissues, hierarchically clustered). Brain tissues cluster separately with elevated ganglioside-related genes. **(B)** PCA biplot (PC1 vs PC2) of tissue-level glycogene expression profiles. Arrows indicate top gene loadings. **(C)** PC1 vs PC3, capturing additional variance along the HS/N-glycan axis. **(D)** Scree plot showing proportion of variance explained.

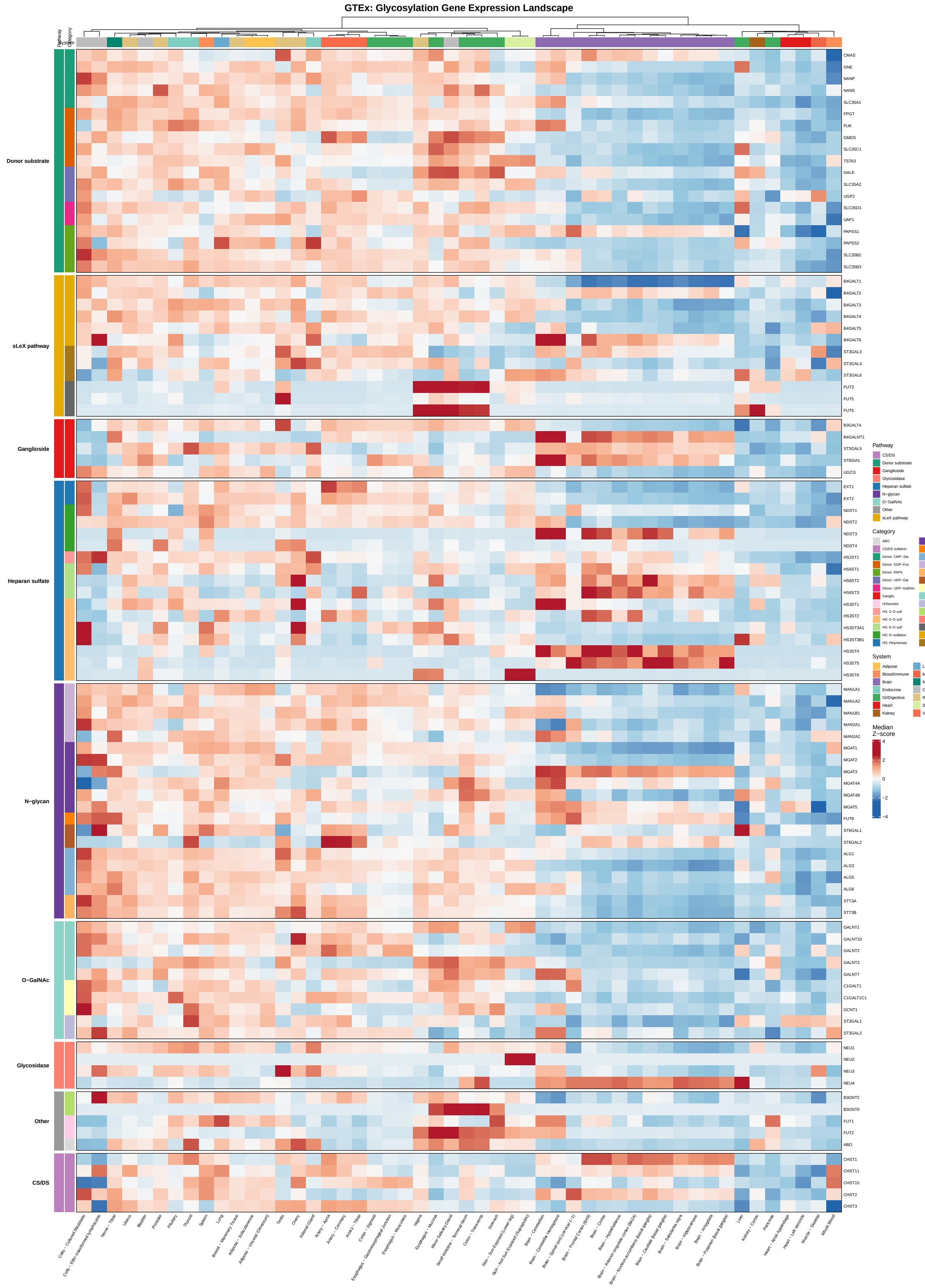


**S2 Fig. Full glycogene expression landscape.** Heatmap of median Z-scored expression for all 97 glycogenes (rows) across 50 GTEx tissues (columns). Genes are grouped by functional category (color sidebar); tissues are hierarchically clustered. Generated using ComplexHeatmap.

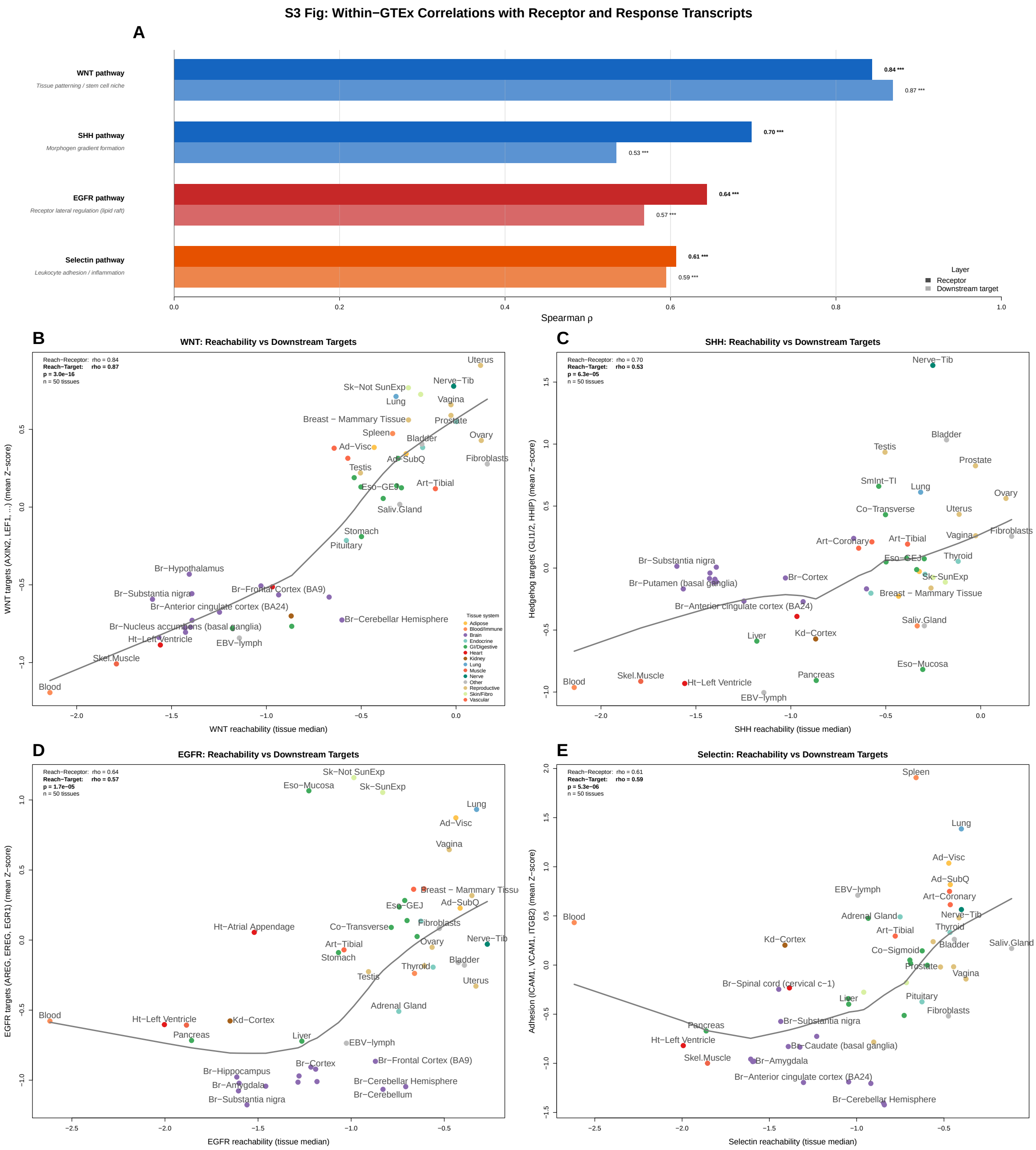


**S3 Fig. Within-GTEx correlations with receptor and pathway-response transcripts.** **(A)** Paired barplot of Spearman $\rho$ for each cascade: reachability vs receptor (dark bars) and reachability vs downstream targets (light bars). **(B–E)** Tissue-level reachability versus response-transcript mean Z-score. Both axes derive from the same RNA-seq and do not establish glycan-mediated signaling.

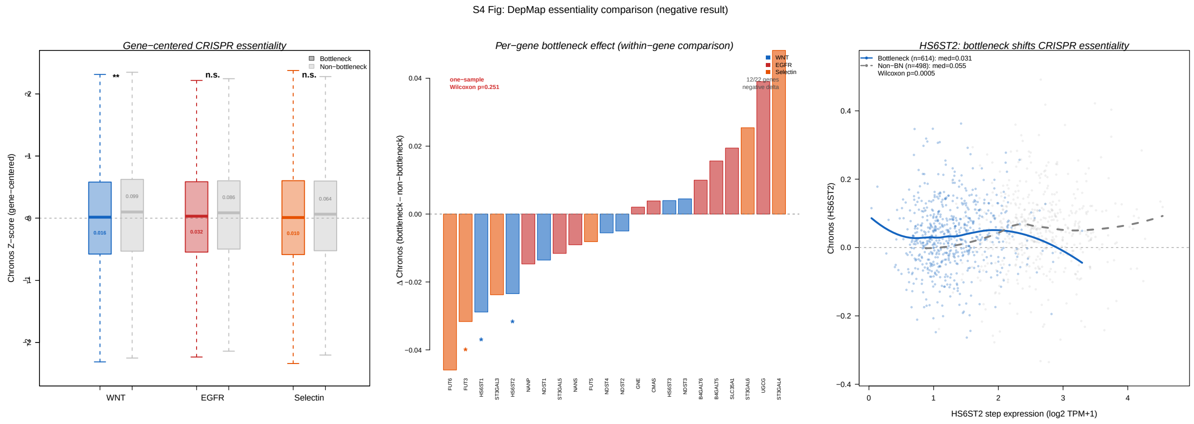

**S4 Fig. DepMap essentiality analysis (negative result).** Comparison of Chronos essentiality scores between cell lines where a glycosylation gene is an expression argmin versus elsewhere, for the three coherence cascades. Only HS6ST1 and HS6ST2 differ after BH correction; the overall result is negative.

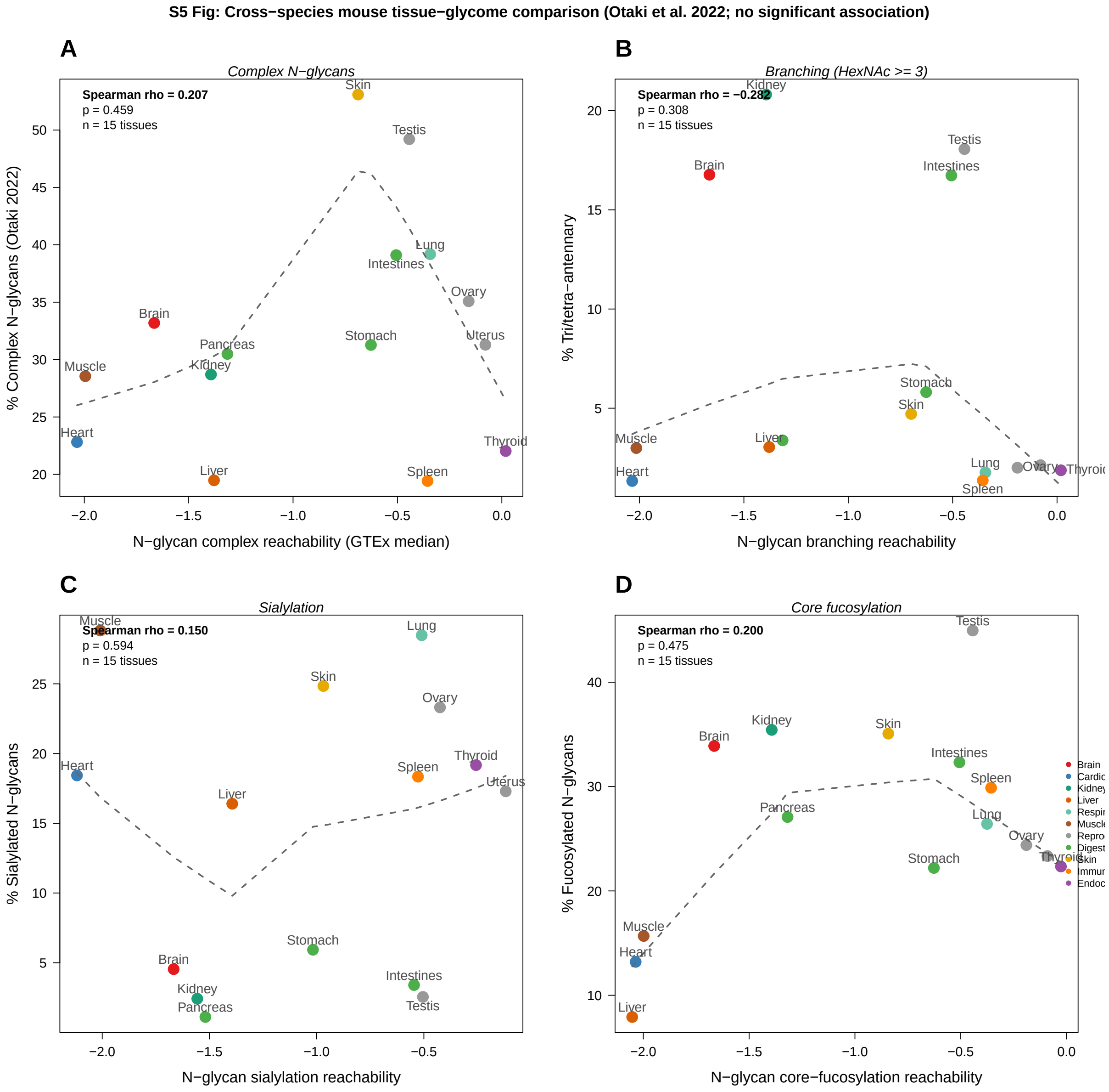


**S5 Fig. Cross-species tissue glycomics comparison (negative result).** Scatter plots of human GTEx-derived N-glycan reachability scores ($x$-axis) versus experimentally measured mouse tissue glycomic features ($y$-axis) from Otaki et al. [2] for $n = 15$ matched tissues, colored by organ system. **(A)** Complex N-glycans ($\rho = 0.207$, $p = 0.459$). **(B)** Tri/tetra-antennary branching ($\rho = -0.282$, $p = 0.308$). **(C)** Sialylation ($\rho = 0.150$, $p = 0.594$). **(D)** Core fucosylation ($\rho = 0.200$, $p = 0.475$). None supports a positive abundance association.

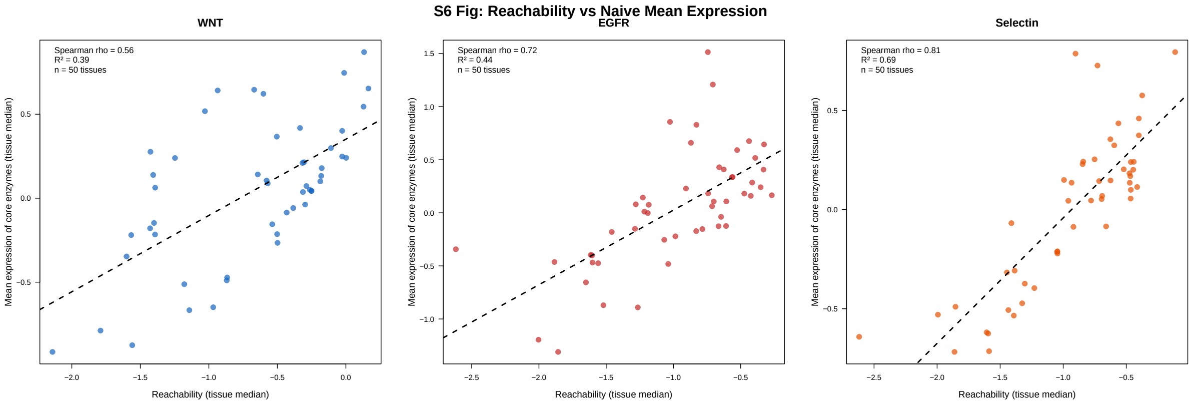


**S6 Fig. Reachability versus naive mean expression.** Scatter plots comparing tissue-median reachability and naive mean expression for the three coherence cascades. Divergence describes different aggregation behavior; it does not identify which score better reflects glycan abundance.

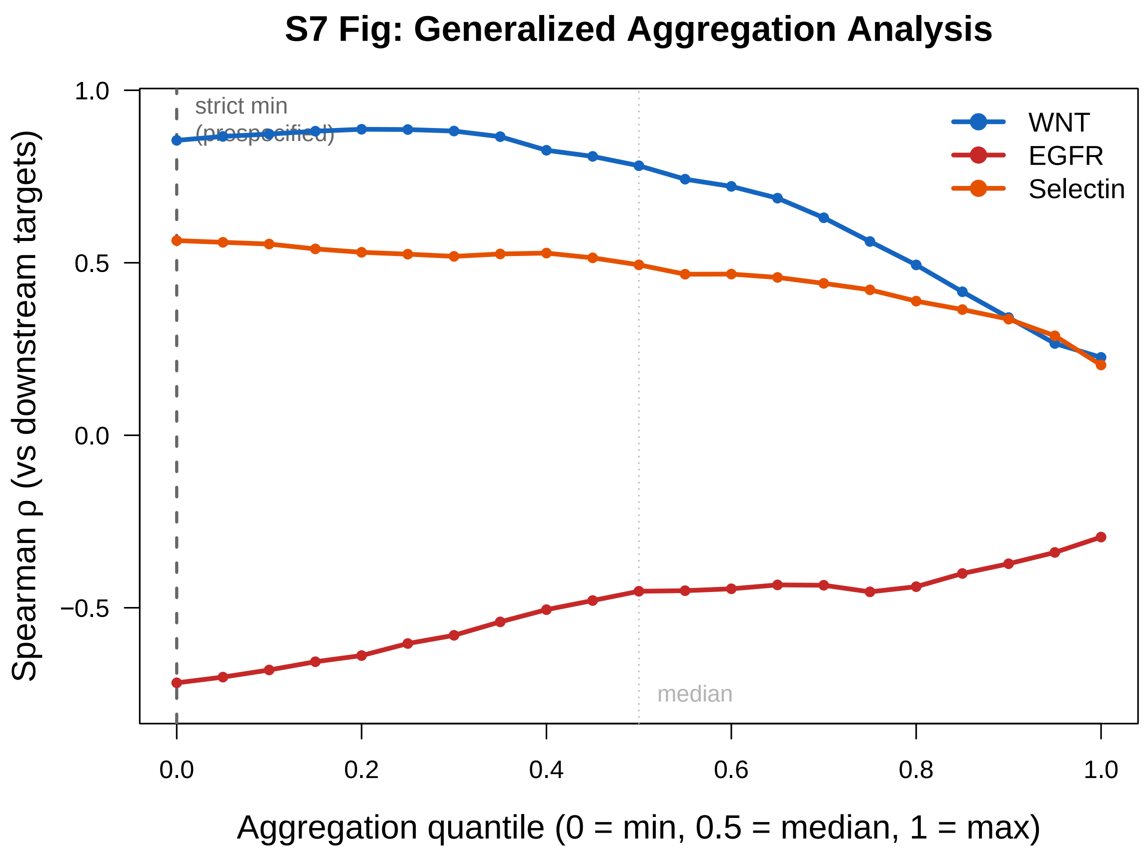


**S7 Fig. AND-aggregation quantile sensitivity.** Within-GTEx direction-oriented response correlation as the AND quantile varies from minimum ($q = 0$) to maximum ($q = 1$), with OR=mean fixed. Negative EGFR values are opposite the prespecified GM3-inhibitory direction. The sweep is descriptive and was not used to select the method.

**S1 Table. Donor substrate demand indices.** Number of reachability metrics requiring each nucleotide sugar donor. CMP-Sia and UDP-Gal are the most widely shared donors (12 and 10 metrics, respectively). Metrics requiring multiple donors (e.g., sLeX requires CMP-Sia, GDP-Fuc, and UDP-Gal) have additional bottleneck opportunities.

| Donor | #Metrics | Example metrics |
|---|---|---|
| CMP-Sia | 12 | sLeX, GM3, GD3, Ng_sia, OGN_sia |
| UDP-Gal | 10 | sLeX, GM1, Ng_complex, Ng_branch, OGN_Core1 |
| GDP-Fuc | 4 | sLeX, Ng_coreFuc, reach_FGF/WNT/SHH |
| PAPS | 8 | HS_N, HS_2O, HS_6O, HS_3O, reach_FGF/WNT/SHH |
| UDP-GalNAc | 6 | GM2, OGN_Tn, OGN_Core1, OGN_Core2, OGN_sia |

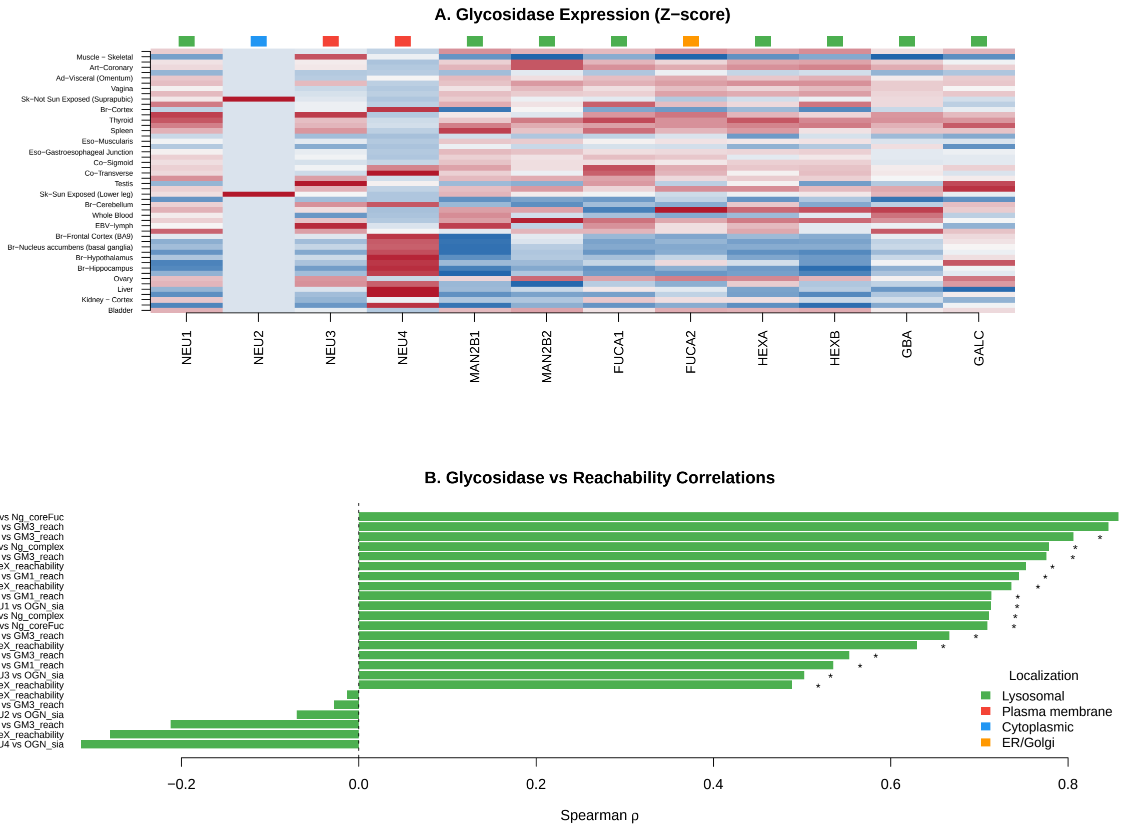


**S8 Fig. Glycosidase expression survey.** Spearman correlations between glycosidase expression and prespecified reachability metrics across 50 tissues. Examples are NEU1 versus GM3 ($\rho = 0.775$) and MAN2B1 versus Ng complex ($\rho = 0.711$). These shared-transcriptome associations neither measure turnover nor remove catabolic confounding.

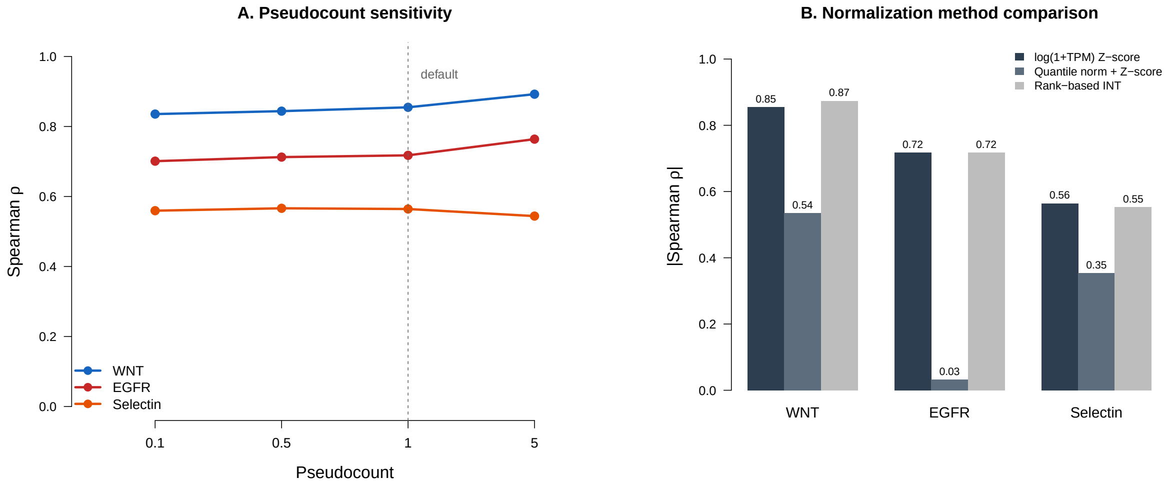


**S9 Fig. Normalization and pseudocount sensitivity.** Direction-oriented coherence correlations under four pseudocounts (0.1, 0.5, 1, 5), quantile normalization, and rank-based inverse-normal transformation. Negative EGFR values are opposite the prespecified GM3-inhibitory direction. Rank normalization is similar to the default, whereas quantile normalization substantially changes the correlations.

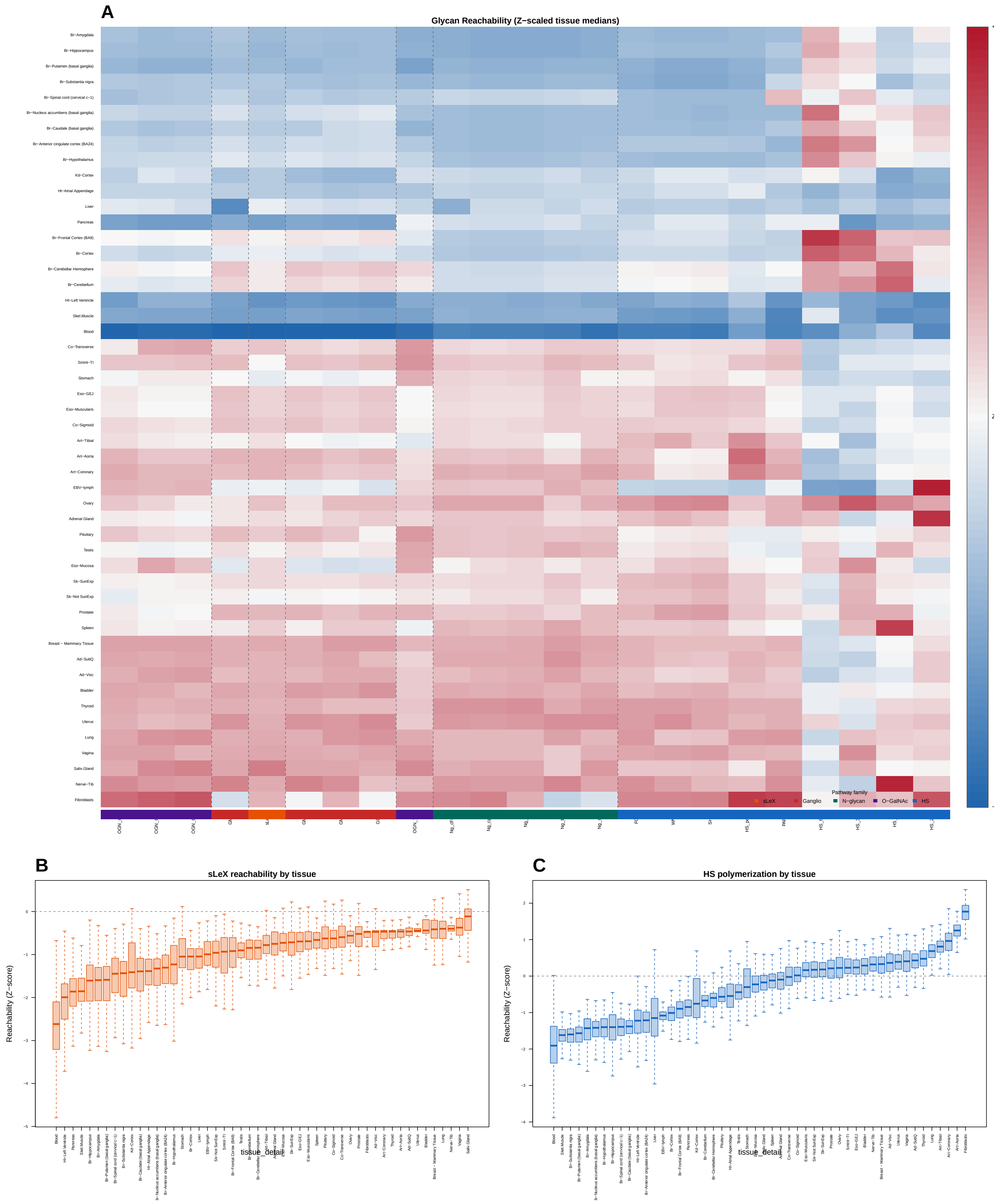


**S10 Fig. Tissue median reachability landscape.** Heatmap of Z-scaled tissue median reachability scores for all 23 metrics (rows) across 50 GTEx tissues (columns). Both axes are hierarchically clustered; tissue axis orientation and abbreviated labels match S1 Fig A for direct comparison. Brain regions form a tight cluster with elevated ganglioside (GM3, GD3) but depressed N-glycan and HS reachability. Connective tissue-rich organs (nerve, fibroblasts, ovary) show the highest HS-related reachability. Whole blood, skeletal muscle, and heart rank lowest across most metrics, reflecting reduced glycosylation machinery in terminally differentiated cells. This heatmap complements the raw expression landscape (S1 Fig A; S2 Fig) by revealing which biosynthetic pathways have expression-complete profiles rather than which individual genes are transcribed.

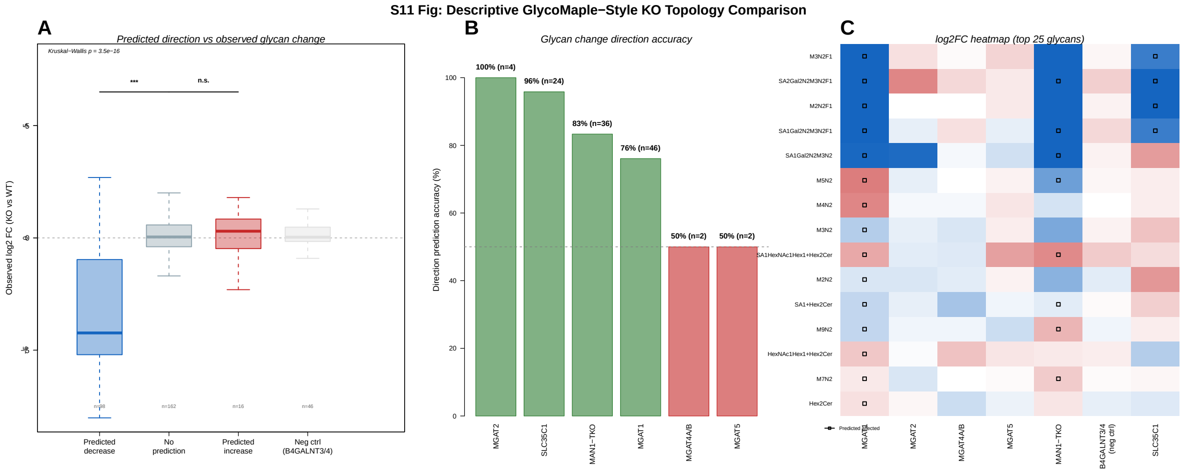


**S11 Fig. Descriptive per-knockout topology detail.** (**A**) Direction-prediction accuracy for six N-glycan-processing knockouts using HEK293 N-glycan mass-spectrometry data [1]. Bar height indicates the percentage of glycan species with correctly predicted direction of change. Early-pathway knockouts (SLC35C1, MGAT2, MGAT1, MAN1A1/A2/B1 triple) show high accuracy; late-pathway knockouts (MGAT4A/B, MGAT5) have few testable glycans. (**B**) Step expression rank versus glycomic perturbation magnitude. These panels are descriptive; the baseline-controlled benchmark is Fig 7, where the minimum does not outperform the mean or binary scorer.

**S2 Table. Summary of within-GTEx coherence sensitivities.** These statistics assess RNA-level association robustness, not glycan-mediated causation. Values are direction-oriented: the EGFR response was multiplied by −1 so that positive would match the prespecified GM3-inhibitory direction; negative EGFR values therefore point in the opposite direction. Complete outputs are in Supplementary File S2.

| Analysis | WNT | EGFR | Selectin | Concern addressed |
|---|---|---|---|---|
| Primary $\rho$ | 0.855 | −0.718 | 0.564 | Tissue-level coherence |
| Donor-bootstrap 95% CI | [0.852, 0.873] | [−0.742, −0.700] | [0.536, 0.581] | Shared donors |
| Variance-matched perm. $p$ | 0.002 | 0.395 | 0.783 | Generic tissue covariance |
| Within-tissue median $\rho$ | 0.549 | −0.385 | 0.354 | Tissue identity |
| Partial $\rho$ (core mean) | 0.795 | −0.743 | 0.508 | Conditional association |
| Partial $\rho$ (all-pathway mean) | 0.680 | −0.641 | 0.387 | Stricter expression control |

**S3 Table. GTEx v8 tissue sample sizes.** The table lists all 54 tissue types. Primary analyses were restricted to the 50 tissues with $N \geq 20$; Cervix Ectocervix, Cervix Endocervix, Fallopian Tube, and Kidney Medulla were excluded.

| Tissue | $N$ | Tissue | $N$ |
|---|---|---|---|
| Adipose - Subcutaneous | 663 | Liver | 226 |
| Adipose - Visceral | 541 | Lung | 578 |
| Adrenal Gland | 258 | Minor Salivary Gland | 162 |
| Artery - Aorta | 432 | Muscle - Skeletal | 803 |
| Artery - Coronary | 240 | Nerve - Tibial | 619 |
| Artery - Tibial | 663 | Ovary | 180 |
| Bladder | 21 | Pancreas | 328 |
| Brain - Amygdala | 152 | Pituitary | 283 |
| Brain - Ant. cing. cortex | 176 | Prostate | 245 |
| Brain - Caudate | 246 | Skin - Not Sun Exp | 604 |
| Brain - Cb. Hemisphere | 215 | Skin - Sun Exposed | 701 |
| Brain - Cerebellum | 241 | Small Intestine | 187 |
| Brain - Cortex | 255 | Spleen | 241 |
| Brain - Frontal Cortex | 209 | Stomach | 359 |
| Brain - Hippocampus | 197 | Testis | 361 |
| Brain - Hypothalamus | 202 | Thyroid | 653 |
| Brain - Nuc. accumbens | 246 | Uterus | 142 |
| Brain - Putamen | 205 | Vagina | 156 |
| Brain - Spinal cord | 159 | Whole Blood | 755 |
| Brain - Subst. nigra | 139 | | |
| Breast | 459 | | |
| Fibroblasts | 504 | | |
| Cervix - Ectocervix | 9 | | |
| Cervix - Endocervix | 10 | | |
| Colon - Sigmoid | 373 | | |
| Colon - Transverse | 406 | | |
| EBV-lymphocytes | 174 | | |
| Esophagus - GEJ | 375 | | |
| Esophagus - Mucosa | 555 | | |
| Esophagus - Muscularis | 515 | | |
| Fallopian Tube | 9 | | |
| Heart - Atrial App. | 429 | | |
| Heart - Left Ventricle | 432 | | |
| Kidney - Cortex | 85 | | |
| Kidney - Medulla | 4 | | |

**S1 File. Machine-readable pathway definitions.** Complete definitions of the five pathway families, 23 metrics, donor-supply chains, and OR=mean/AND=min rules in XLSX format.

**S2 File. Complete benchmark and sensitivity analyses.** Workbook containing KO benchmark pairs and summaries, nested quantile selection, all 950 tissue–metric bottleneck-stability rows, ST6GAL sensitivity results, and multiple-testing families.

### References


[1] Huang YF, Aoki K, Akase S, Ishihara M, Liu YS, Yang G, et al. Global mapping of glycosylation pathways in human-derived cells. *Dev Cell.* 2021;56(8):1195–1209.

[2] Otaki M, Hirane N, Natsume-Kitatani Y, Nogami Itoh M, Shindo M, Kurebayashi Y, et al.

**S4 Table. Master list of all 23 reachability metrics.** Each metric represents a distinct endpoint or pathway module. All 23 metrics are included in the pairwise tissue analysis; the Type column preserves the original endpoint/composite classification. Code variables match the analysis outputs.

| Code variable | Display name | Pathway family | Type | #Steps | #Genes |
|---|---|---|---|---|---|
| sLeX_reachability | sLeX reachability | Sialyl Lewis X | Primary | 3 | 12 |
| GM3_reach | GM3 reach | Ganglioside | Primary | 3 | 4 |
| GM2_reach | GM2 reach | Ganglioside | Primary | 4 | 5 |
| GM1_reach | GM1 reach | Ganglioside | Primary | 5 | 6 |
| GD3_reach | GD3 reach | Ganglioside | Primary | 4 | 5 |
| HS_poly | HS polymerization | Heparan sulfate | Primary | 1 | 2 |
| PAPS | PAPS | Donor supply | Primary | 2 | 4 |
| HS_N | HS N-sulfation | Heparan sulfate | Primary | 2 | 6 |
| HS_2O | HS 2-O sulfation | Heparan sulfate | Primary | 2 | 3 |
| HS_6O | HS 6-O sulfation | Heparan sulfate | Primary | 2 | 5 |
| HS_3O | HS 3-O sulfation | Heparan sulfate | Primary | 2 | 9 |
| reach_FGF_like | HS FGF-like | HS composite | Composite | 4 | 11 |
| reach_WNT_like | HS WNT-like | HS composite | Composite | 4 | 11 |
| reach_SHH_like | HS SHH-like | HS composite | Composite | 5 | 20 |
| Ng_complex | Ng complex | N-glycan | Composite | 7 | 21 |
| Ng_branch | Ng branch | N-glycan | Composite | 8 | 24 |
| Ng_coreFuc | Ng core Fuc | N-glycan | Composite | 8 | 22 |
| Ng_bisect | Ng bisect | N-glycan | Composite | 8 | 22 |
| Ng_sia | Ng sialylation | N-glycan | Composite | 8 | 23 |
| OGN_Tn | OGN Tn | O-GalNAc | Composite | 1 | 20 |
| OGN_Core1 | OGN Core 1 | O-GalNAc | Composite | 2 | 22 |
| OGN_Core2 | OGN Core 2 | O-GalNAc | Composite | 3 | 23 |
| OGN_sia | OGN sialylation | O-GalNAc | Composite | 3 | 24 |

**S5 Table. Tissue name abbreviations used in figures.** Mapping between full GTEx v8 tissue names and abbreviated labels used in heatmaps and scatter plots throughout this study. Tissues whose full name is used without abbreviation (e.g., Liver, Lung, Ovary) are omitted. $N$: number of RNA-seq samples per tissue.

| GTEx tissue name | Abbreviation | $N$ |
|---|---|---|
| Adipose - Subcutaneous | Ad-SubQ | 663 |
| Adipose - Visceral (Omentum) | Ad-Visc | 541 |
| Adrenal Gland | Adrenal | 258 |
| Artery - Aorta | Art-Aorta | 432 |
| Artery - Coronary | Art-Coronary | 240 |
| Artery - Tibial | Art-Tibial | 663 |
| Brain - Amygdala | Br-Amygdala | 152 |
| Brain - Anterior cingulate cortex (BA24) | Br-ACC | 176 |
| Brain - Caudate (basal ganglia) | Br-Caudate | 246 |
| Brain - Cerebellar Hemisphere | Br-CbHemi | 215 |
| Brain - Cerebellum | Br-Cerebellum | 241 |
| Brain - Cortex | Br-Cortex | 255 |
| Brain - Frontal Cortex (BA9) | Br-FrCortex | 209 |
| Brain - Hippocampus | Br-Hippocampus | 197 |
| Brain - Hypothalamus | Br-Hypothalamus | 202 |
| Brain - Nucleus accumbens (basal ganglia) | Br-NAcc | 246 |
| Brain - Putamen (basal ganglia) | Br-Putamen | 205 |
| Brain - Spinal cord (cervical c-1) | Br-SpCord | 159 |
| Brain - Substantia nigra | Br-SN | 139 |
| Breast - Mammary Tissue | Breast | 459 |
| Cells - Cultured fibroblasts | Fibroblasts | 504 |
| Cells - EBV-transformed lymphocytes | EBV-lymph | 174 |
| Colon - Sigmoid | Co-Sigmoid | 373 |
| Colon - Transverse | Co-Transverse | 406 |
| Esophagus - Gastroesophageal Junction | Eso-GEJ | 375 |
| Esophagus - Mucosa | Eso-Mucosa | 555 |
| Esophagus - Muscularis | Eso-Muscularis | 515 |
| Heart - Atrial Appendage | Ht-Atrial Appendage | 429 |
| Heart - Left Ventricle | Ht-Left Ventricle | 432 |
| Kidney - Cortex | Kd-Cortex | 85 |
| Kidney - Medulla | Kd-Medulla | 4 |
| Minor Salivary Gland | Saliv.Gland | 162 |
| Muscle - Skeletal | Skel.Muscle | 803 |
| Nerve - Tibial | Nerve-Tib | 619 |
| Skin - Not Sun Exposed (Suprapubic) | Sk-NoSun | 604 |
| Skin - Sun Exposed (Lower leg) | Sk-SunExp | 701 |
| Small Intestine - Terminal Ileum | SmInt-TI | 187 |
| Whole Blood | Blood | 755 |

Mouse tissue glycome atlas 2022 highlights inter-organ variation in major N-glycan profiles. *Sci Rep*. 2022;12(1):17804.
